# RainbowSTORM: An open-source ImageJ plugin for spectroscopic single-molecule localization microscopy (sSMLM) data analysis and image reconstruction

**DOI:** 10.1101/2020.03.10.986018

**Authors:** Janel L. Davis, Brian Soetikno, Ki-Hee Song, Yang Zhang, Cheng Sun, Hao F. Zhang

## Abstract

**Summary:** Spectroscopic single-molecule localization microscopy (sSMLM) simultaneously captures the spatial locations and full spectra of stochastically emitting fluorescent single molecules. It provides an optical platform to develop new multi-molecular and functional imaging capabilities. While several open-source software suites provide sub-diffraction localization of fluorescent molecules, software suites for spectroscopic analysis of sSMLM data remain unavailable. RainbowSTORM is an open-source, user-friendly ImageJ/FIJI plugin for end-to-end spectroscopic analysis and visualization for sSMLM images. RainbowSTORM allows users to calibrate, preview, and quantitatively analyze emission spectra acquired using different reported sSMLM system designs and fluorescent labels.

**Availability:** RainbowSTORM is a java plugin for ImageJ (https://imagej.net)/FIJI (http://fiji.sc) freely available through: https://github.com/FOIL-NU/RainbowSTORM. RainbowSTORM has been tested with Windows and Mac operating systems and ImageJ/FIJI version 1.52.

**Supplementary information:** Supplementary data are available online.

## 1. Introduction

Single-molecule localization microscopy (SMLM) (Betzig, et al., 2006; Rust, et al., 2006; Sharonov and Hochstrasser, 2006) overcomes the optical diffraction limit by localizing stochastically emitting fluorescent molecules with high localization precision (typically 10-20 nm). Recently, spectroscopic single-molecule localization microscopy (sSMLM) (Bongiovanni, et al., 2016; Dong, et al., 2016; Zhang, et al., 2015), which simultaneously detects the location and full emission spectra of each emission event was reported. Thus far, sSMLM has enabled multi-color imaging (Zhang, et al., 2015) and tracking (Huang, et al., 2018) of as many as four different fluorescent species using a single excitation source. sSMLM has also led to new functional imaging capabilities through analyzing variations in the spectra of individual molecules. For example, sSMLM detected the polarity of the environment surrounding dye molecules (Bongiovanni, et al., 2016) and enabled the discovery of previously undetected molecular conformations of dyes (Kim, et al., 2017). Overall, sSMLM shows great promise to further extend existing SMLM. While a variety of software algorithms and packages are currently available for processing and analyzing traditional SMLM images (Sage, et al., 2019), software tools for comprehensive spectroscopic analysis of sSMLM images remain unavailable.

Here, we present RainbowSTORM, an open-source spectroscopic analysis plugin for ImageJ/FIJI. RainbowSTORM leverages the functionality of the existing SMLM processing tool Thunder-STORM (Ovesny, et al., 2014) to attain spatial information while providing crucial spectroscopic tools for system calibration as well as spectral identification and classification. RainbowSTORM uses the spectral centroids (or intensity-weighted spectral means) of each localized stochastic event to define a range of spectral colors and render pseudo-colored sSMLM super-resolution images (Bongiovanni, et al., 2016; Dong, et al., 2016; Zhang, et al., 2015). Multicolor images can be generated by setting different user-defined spectral centroid ranges for channels with predefined colors. We provide test calibration and sSMLM images along with a testing protocol and a detailed user guide which includes descriptions and workflows for the processes implemented in RainbowSTORM. Derivations for spectroscopic analysis (Song, et al., 2018) and flowcharts of the algorithms used in RainbowSTORM are included in the supplementary information.

## 2. Features and Methods

### 2.1 System calibration

RainbowSTORM calibrates sSMLM images acquired using systems, where the dispersive element (Figure 1a) can be either a grating or a prism. While grating-based systems are calibrated by linearly fitting pixel positions to known wavelengths (Dong, et al., 2016), prism-based systems are calibrated using second-order (Huang, et al., 2018) or third-order (Zhang, et al., 2015) polynomial fittings. Calibration in RainbowSTORM can be performed using both calibrated light sources (e.g. calibration lamps or multiple laser lines) and multi-color fluorescent beads.

**Figure 1.**
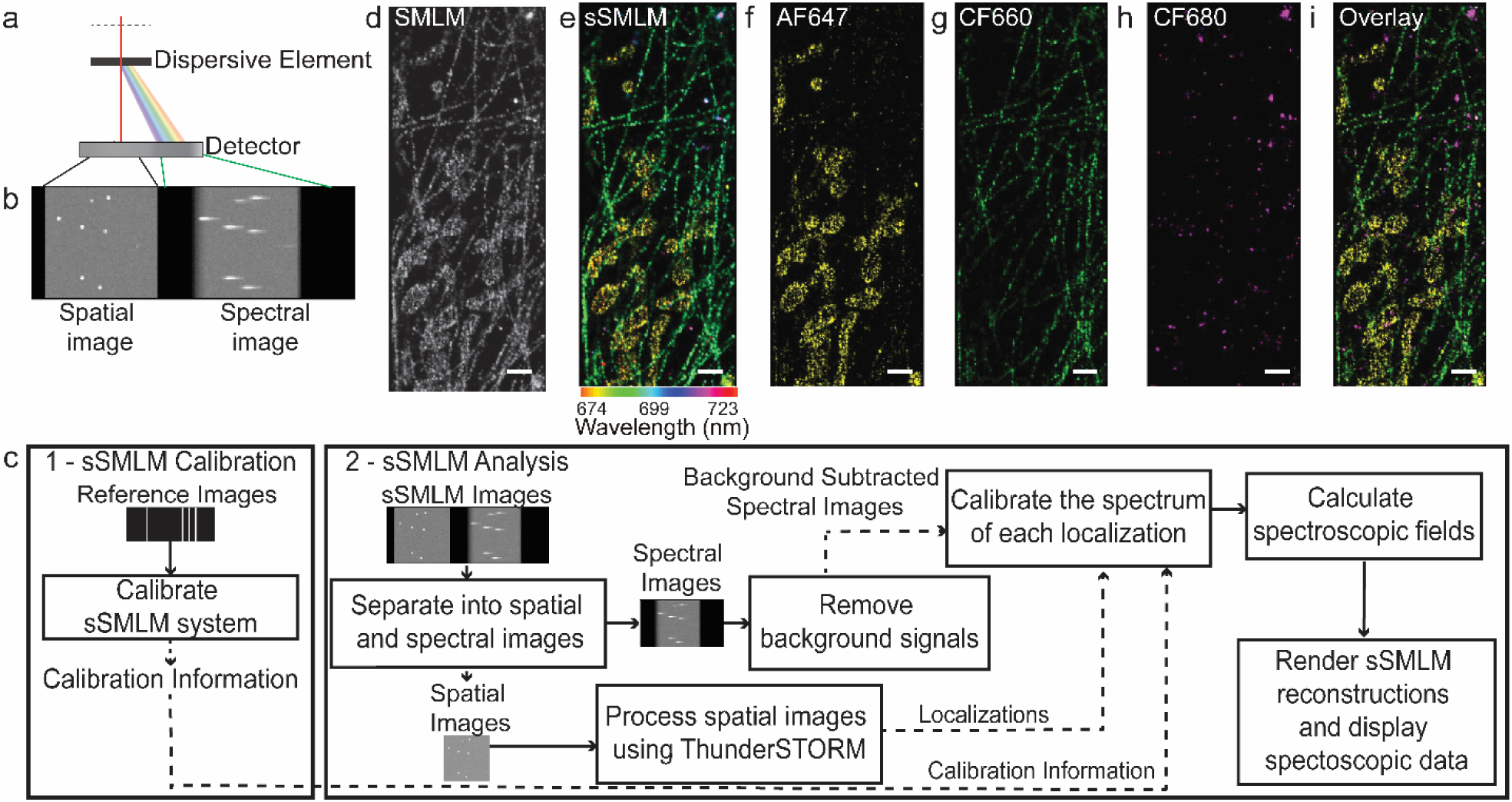
(a) General sSMLM system schematic. (b) sSMLM images with the spatial and spectral images simultaneously captured on different parts of a detector. (c) RainbowSTORM workflow showing how the system calibration module interacts with the analysis module (d) SMLM reconstruction (e) Pseudo-colored sSMLM reconstruction. Images of the three separate channels showing (f) mitochondria labeled with AF647, (g) microtubules labeled with CF660, (h) peroxisomes labeled with CF680, and (i) the overlay image of the three channels.

### 2.2 sSMLM image processing

In addition to providing a flexible calibration tool, RainbowSTORM also includes a sSMLM analysis module for processing sSMLM images (Figure 1b). The general workflow for RainbowSTORM analysis is outlined in Figure 1c. RainbowSTORM first requires sSMLM images to be cropped for spatial and spectral analysis. Next two-dimensional (2D) spatial images and three-dimensional (3D) spatial images, captured using the astigmatism method (Huang, et al., 2008; Zhang, et al., 2015), can be processed using ThunderSTORM. Figure 1d shows the resulting SMLM reconstruction after spatial analysis. Next, RainbowSTORM can be used to remove background signals from the spectral images, automatically exclude emission events which spatially overlap, and preview results of spectroscopic analysis using the current processing parameters. Finally, RainbowSTORM identifies the full spectra and calculates the spectroscopic fields for all localizations.

### 2.3 Visualization and post-processing

After processing the spectral images, pseudo-colored super-resolution reconstructions (Figure 1e) are rendered using the spectral centroids and spatial coordinates of each localization. For 3D sSMLM images, a stack of pseudo-colored super-resolution reconstructions can be rendered, where images in the stack are separated by the axial position of each localization. The histograms of the calculated spatial and spectral fields for the processed localizations can be displayed and used to select subsets of the data for independent visualization. Additionally, localizations with large point spread function and spectrum widths as well as localizations with low photon counts and precisions in the spatial and spectral domains can be excluded. RainbowSTORM can also apply ThunderSTORM drift-correction files and assess the image quality of sSMLM images using Fourier Ring Correlation (FRC) analysis (Nieuwenhuizen, et al., 2013). In addition, the spectral centroid information can be assigned to multiple channels to create multicolor super-resolution images using the classification module. For example, Figures 1d-h shows images of the mitochondria, microtubules, and peroxisomes of COS-7 cells respectively labeled by Alexa Fluor 647(AF647), CF660, and CF680. Figure 1i shows the overlay of the three images from the selected spectral centroid windows. After post-processing, sSMLM results can be saved. Previously saved sSMLM results can be loaded using the sSMLM import module for further analysis.

## 3. Summary

RainbowSTORM provides a plugin for performing spectroscopic analysis of 2D and 3D sSMLM images acquired using both grating-based and prism-based sSMLM implementations. RainbowSTORM fills the need for a spectroscopic analysis platform and provides spectral classification methods, spectral and spatial filtering methods, pseudo-colored visualization of sSMLM datasets, and built-in FRC analysis. Future updates will make RainbowSTORM compatible with a wider range of spatial analysis platforms. Further, additional spectroscopic analysis methods such as spectral unmixing (Davis, et al., 2018), machine-learning based spectral classification (Zhang, et al., 2019), and cluster analysis (Bongiovanni, et al., 2016) will be added to RainbowSTORM.

## Supporting information

RainbowSTORM Tutorial

Calibration Images

Simulated sSMLM Images

RainbowSTORM User Guide

