## Supplementary material for "RainbowSTORM: An open-source ImageJ plugin for spectroscopic single-molecule localization microscopy (sSMLM) data analysis and image reconstruction": RainbowSTORM Tutorial

1. Install RainbowSTORM

   i. Download the RainbowSTORM plugin (Rainbow_STORM.jar)
   ii. Copy the file into the Plugins subfolder of your ImageJ installation (e.g. "C:\Program Files\ImageJ\plugins")
   iii. Verify the successful installation of RainbowSTORM
   iv. Restart ImageJ
   v. Locate RainbowSTORM under the Plugins menu
   vi. Download and install the latest version of ThunderSTORM
   vii. For ImageJ installations the Bioformats plugin is also needed to load the provided test datasets.

2. Calibrate the System
   i. Download the calibration images (Calibration.tif)
      - Calibration Image Properties;
        - Calibration Source: Calibration Lamp
        - Dispersive Element: Grating
   ii. Open ImageJ
   iii. Drag the calibration images into the ImageJ window or load the calibration images
   iv. Launch the RainbowSTORM Calibration Screen (from the ImageJ menu Plugins→RainbowSTORM→sSMLM Calibration)

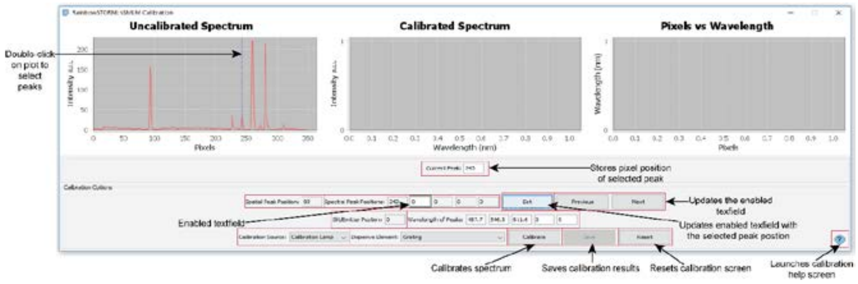


Figure 1: RainbowSTORM Calibration Screen

   v. Double-click on the line plot displayed in the uncalibrated spectrum graph to update the value of the current peak value text field starting with the first peak representing the spatial peak position). Click and drag to zoom in on an area.
   vi. Press the 'Set' button to update the activated text field. Use the 'Previous' and 'Next' buttons to change which text field is activated.
   vii. Select the spectral peaks for example the default wavelengths correspond to the spectral peaks at 487.7 nm, 546.5 nm and 611.6 nm indicated in Figure 1 below:

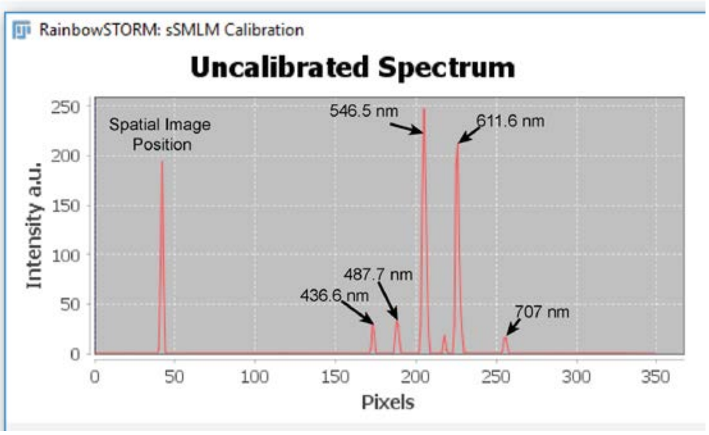


**Figure 2:** Example of spatial and spectral peaks for the example calibration image

viii. The default settings for the calibration source (calibration light source) and the dispersive element(grating) are appropriate for these calibration images

ix. Press the 'Calibrate' button. The R-squared and RMSE values on the pixel versus wavelength plot to assess the calibration results.

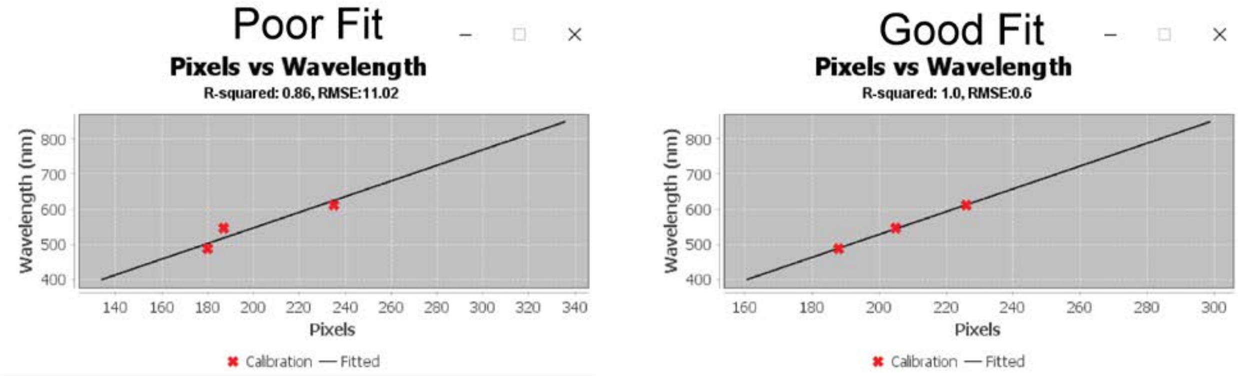


**Figure 3:** Example of poor versus good fitting of calibration results

x. Save the resulting pixel positions, corresponding wavelength information, and fitting method

xi. Close the RainbowSTORM Calibration screen

3. Analyze the sSMLM Images

   i. Download the sSMLM images (Test_A_stack_0107.tif)

   ii. Drag the sSMLM images into the ImageJ window or load the sSMLM images

   iii. Launch the RainbowSTORM sSMLM Analysis Screen (Plugins→ RainbowSTORM→sSMLM Analysis)

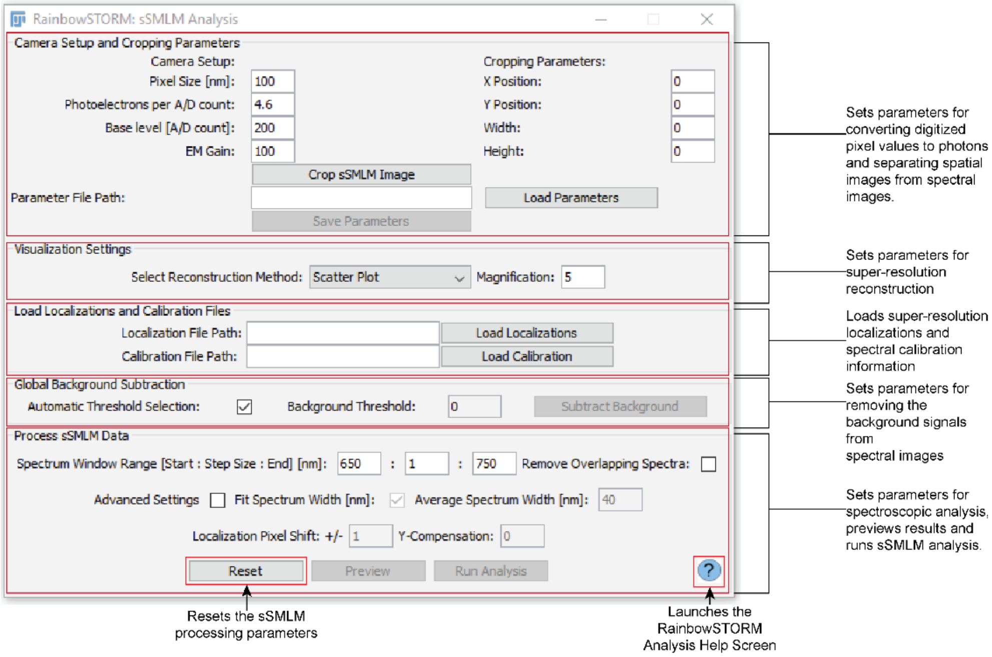


**Figure 4**: RainbowSTORM Analysis Screen

iv. Set the camera parameters

- Camera pixel size: 160 nm
- Photoelectrons per A/D count: 4.6
- Base level [A/D counts]: 200
- EM Gain: 100

v. Use the Rectangular Selection tool from ImageJ's toolbar to select the spatial image in the sSMLM image (see Figure 5) or manually input the parameters of the rectangle into the text fields (initial x position, initial y position, rectangle width, and rectangle height) on the sSMLM Analysis screen

vi. Press the 'Crop sSMLM Image' button on the sSMLM Analysis screen to separate the sSMLM images into spatial and spectral images

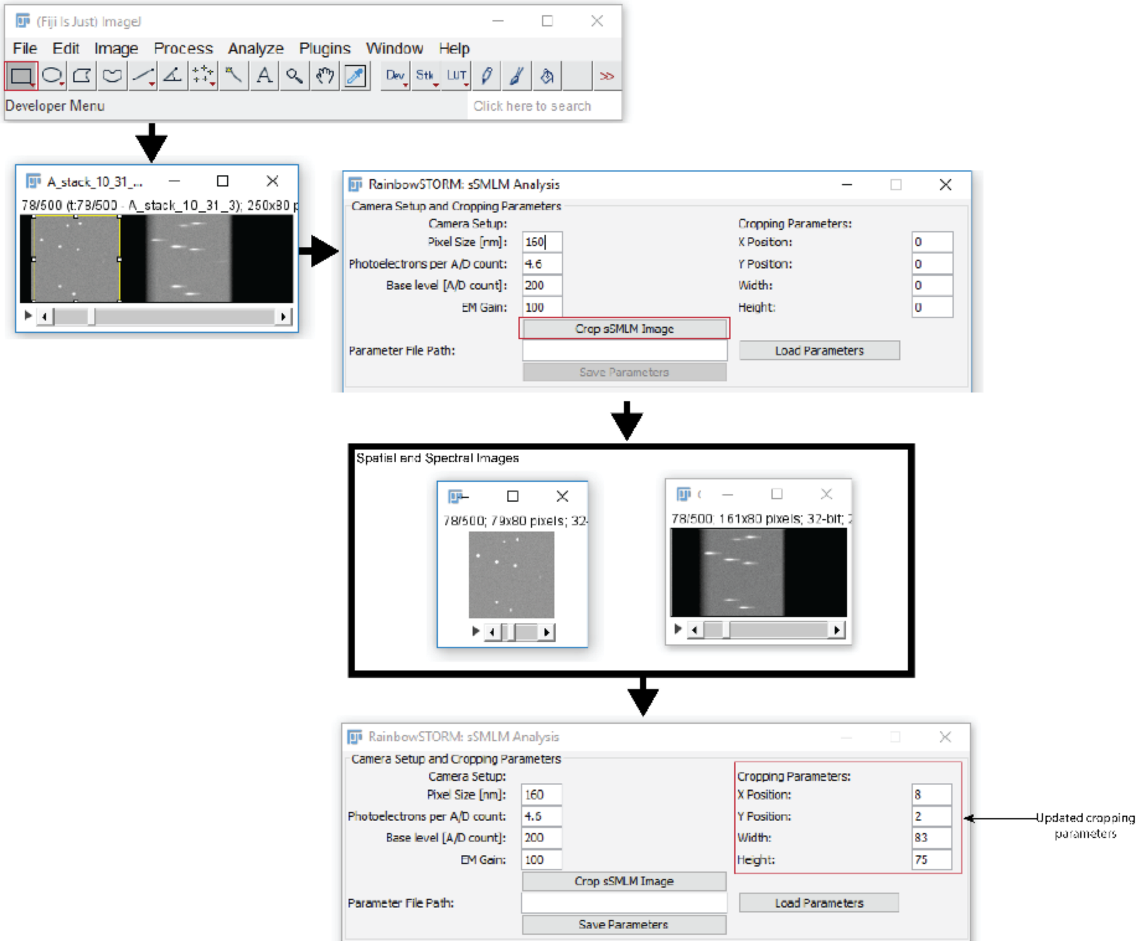


**Figure 5**: Example of the separating the spatial images from the spectra images

vii. Optional-Press the 'Save Parameters' button to save the camera setup and cropping parameters (use an appropriate name to prevent overwriting other RainbowSTORM files)

viii. Select the Spatial Region (Cropped Region 1) and open ThunderSTORM (Plugins→ ThunderSTORM→Run Analysis)

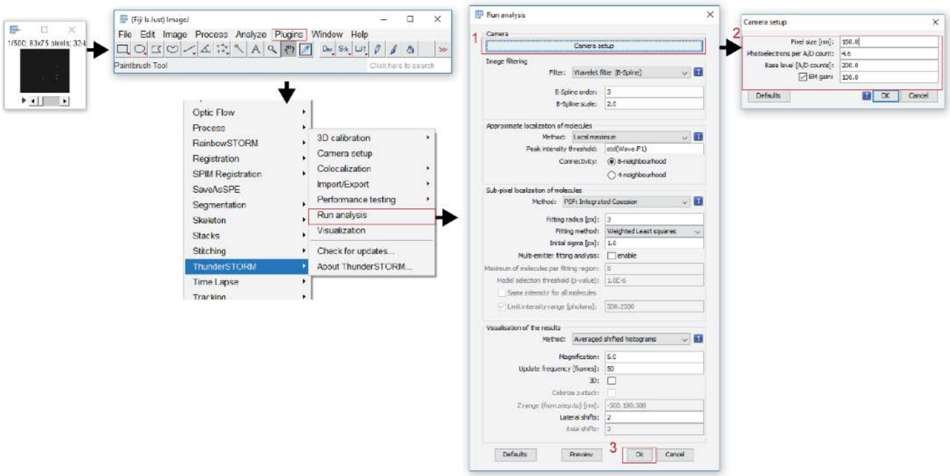


**Figure 6**: Example of analyzing the spatial images using ThunderSTORM

ix. Press the Camera Setup button. Make sure the camera settings used in ThunderSTORM and RainbowSTORM are the same. Press 'OK'.

x. Press 'Ok' to analyze the spatial images using ThunderSTORM (all other default settings are appropriate).

xi. Export the ThunderSTORM localization results (don't overwrite any of the previously saved files)

- Optional- deselect unwanted fields (Required ThunderSTORM fields: frame, x, y, intensity, sigma and uncertainty)

xii. Select the drift correction tab and select the cross-correlation default options. Check the 'Save to file' checkbox and press the ellipses '…' button to specify the filename. Press the 'Apply' button to apply and save the drift correction.

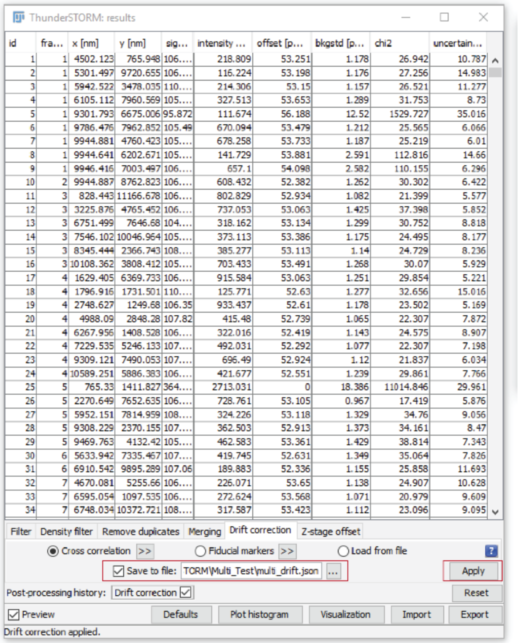


**Figure 7**: Example of saving ThunderSTORM drift correction information

xiii. Close ThunderSTORM

xiv. Select the open RainbowSTORM Analysis screen

xv. Using the ‘Load Localization and Calibration Information’ Panel on the Analysis screen, load the ThunderSTORM localization results into RainbowSTORM

xvi. Load the Calibration Information into RainbowSTORM

xvii. With the Automatic Background Subtraction' checkbox, press the 'Subtract' button

xviii. Default settings are optimized for Far Red dyes e.g. Alexa Fluor 647

xix. Optional - Check the 'Remove Overlapping Spectra' checkbox to excluded localizations with overlapping spectra

xx. Optional - Preview the results of the spectroscopic analysis for individual localizations

xxi. Press the 'Run Analysis' to calculate the results of the spectroscopic analysis for all localizations

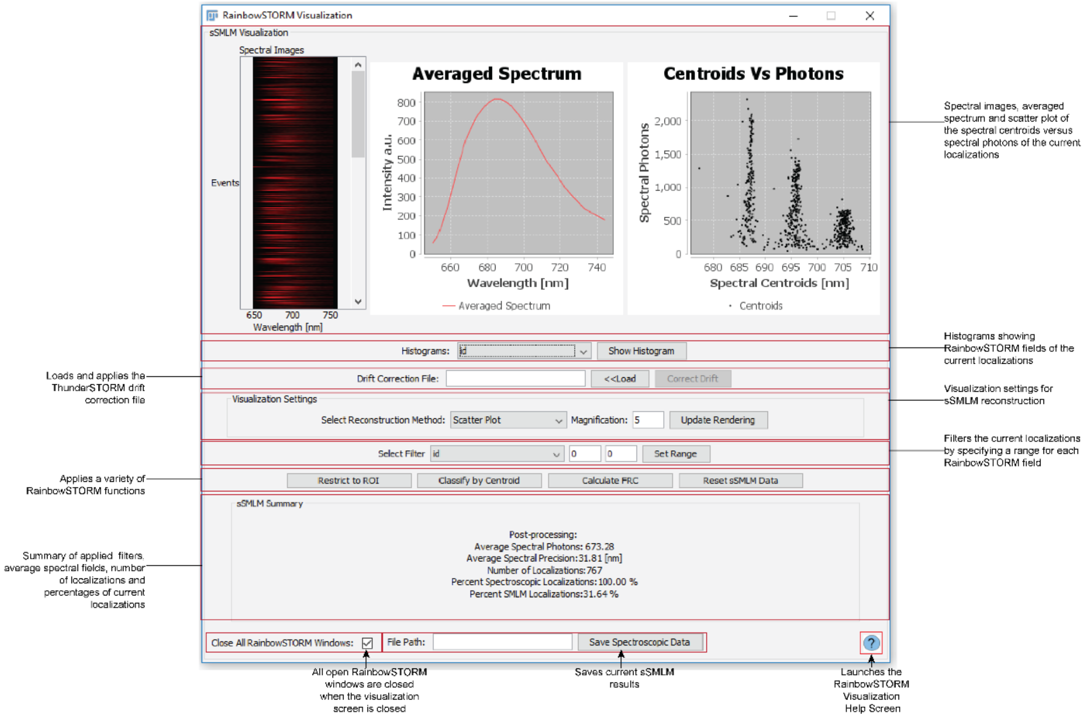


**Figure 8:** RainbowSTORM Visualization Screen

xxii. Select a field (e.g. spectral sigma) from the histograms of the localizations for each RainbowSTORM field can be generated by selecting the field and pressing the "Show Histogram" button.

xxiii. Use the selected histogram to identify and a minimum and maximum value for the Filter Range. Input the values (e.g. spectral sigma 17 nm to 28 nm) and press the 'Set Range' button.

xxiv. Update the Visualization settings:
- Reconstruction method: Averaged Gaussian
- Magnification: 5

xxv. Load the drift correction file and press the correct drift button

xxvi. Press the 'Calculate FRC' to show results from Fourier Ring Correlation (FRC) Analysis

xxvii. Close FRC Results

xxviii. Use the ImageJ's rectangular selection tool is used to select a region of interest (ROI) in the current sSMLM reconstruction then pressing the "Restrict to ROI" button to select localizations within the ROI.

xxix. Press the 'Reset sSMLM Data' button to load original data.

xxx. Press the "Classify by Centroid" button launches the RainbowSTORM Classification screen where localizations can be grouped into color-coded image windows based on their spectral centroids.
- Set any three spectral windows (W1: 680 nm-689 nm, W2: 692 nm-698 nm and W3: 703 nm-709 nm)

- Optional - Check the ‘Show Channel’ checkbox to display a reconstruction of each individual channel

xxxi. Press the ‘Save Spectroscopic Data’ button to save the current spectroscopic data.

xxxii. Close the Visualization screen.

4. Import previously saved data
    i. Launch the Import Screen
        - Select Plugins→ RainbowSTORM→sSMLM Import from the ImageJ menu
    ii. Select the previously saved data ending in “_spec.csv” into RainbowSTORM
    iii. Set Camera Setup parameters to the values used for analysis or load the previously saved camera setup parameters.
    iv. Press the ‘Visualize Data’ button to render the Pseudo-colored image and load the data into the Visualization Screen.
5. Access Help Screens
    i. Launch the RainbowSTORM Help Screen (from the ImageJ menu Plugins→RainbowSTORM→RainbowSTORM Help)
    ii. Use hyperlinks to view help for each RainbowSTORM screen
    iii. Help pages for each screen can be accessed by pressing the blue question mark icon on each screen.
    iv. Close the Help Screens
