## Supplementary material for "RainbowSTORM: An open-source ImageJ plugin for spectroscopic single-molecule localization microscopy (sSMLM) data analysis and image reconstruction": RainbowSTORM User Guide

**Janel L. Davis[1, †], Brian Soetikno[1, †], Ki-Hee Song[1], Yang Zhang[1], Cheng Sun[2] and Hao F. Zhang[1, *]**

[1]Department of Biomedical Engineering, Northwestern University, Evanston, Illinois, USA.
[2]Department of Mechanical Engineering, Northwestern University, Evanston, Illinois USA

† These authors contributed equally

### 1. About RainbowSTORM

RainbowSTORM is an open-source ImageJ/FIJI plugin for analysis of spectroscopic Single Molecule Localization Microscopy (sSMLM) images collected using a single imaging sensor (EMCCD or CMOS). Starting with calibration images recorded using a calibration lamp, fluorescent beads, or both, RainbowSTORM can calibrate both grating-based and prism-based sSMLM systems. The resulting calibration file can be saved and used to process experimental sSMLM images. Using RainbowSTORM's sSMLM cropping module, the recorded sSMLM frames are separated into spatial and spectral images. The spatial images are then analyzed using ThunderSTORM, an open-source ImageJ/FIJI plugin for processing Single Molecule Localization Microscopy (SMLM) images (Ovesny, et al., 2014). The spatial information is then saved and loaded into RainbowSTORM. Next, RainbowSTORM removes the background from the spectral images. Finally, the corresponding spectral image from each single-molecule emission event is identified and calibrated. The resulting spectroscopic information is used to render Pseudo-colored sSMLM super-resolution image reconstructions. Multicolor super-resolution image reconstructions can also be rendered using RainbowSTORM's spectra classification screen.

### 2. Installation

The RainbowSTORM plugin (Rainbow_STORM.jar) can be downloaded from https://github.com/FOIL-NU/RainbowSTORM. To install the plugin, copy the file into the Plugins subfolder of your ImageJ/FIJI installation (e.g. "C:\Program Files\ImageJ\plugins"). To verify successful installation of RainbowSTORM, restart ImageJ/FIJI and locate RainbowSTORM under the Plugins menu. RainbowSTORM also requires ThunderSTORM which can be downloaded from https://github.com/zitmen/thunderstorm/wiki/Downloads. For ImageJ installations the Bioformats plugin (https://www.openmicroscopy.org/bio-formats/downloads/) is also required to load the provided test data.

### 3. System Calibration

RainbowSTORM processes calibration images from both calibration light sources and fluorescent beads. Additionally, RainbowSTORM is compatible with grating-based (Dong, et al., 2016) and prism-based (Zhang, et al., 2015) sSMLM systems. RainbowSTORM's calibration screen can be launched using a single frame, however, it is recommended that at least 100 frames be used.

Upon launching RainbowSTORM's calibration screen (Figures 1 and 2), the background from the calibration images are removed and the averaged background-removed image is generated. However, for samples with uneven backgrounds the Background Subtraction tool under the Process menu can be used prior to launching the calibration screen. Next, the uncalibrated line profile of the average background-removed calibration image is displayed. The user then selects the peak pixel positions from the line profile which correspond to the pixel location of the source in the spatial image and the known wavelength information from the spectral image. By pressing the 'Calibrate' button the system is calibrated, and the user can save the calibration information, which is used on the sSMLM analysis screen to process experimental sSMLM datasets. Examples of calibration using both calibration sources and sSMLM systems are included below.

#### 3.1 Calibration source

The calibration screen is used to relate the spatial information to the spectral information. This can be done using a calibration light source, fluorescent beads, or a combination of them.

##### 3.1.1 Calibration light source

Calibration images can be acquired by a sSMLM system using a calibration lamp or multiple laser lines to illuminate a narrowed slit. The resulting images feature the spatial image of the calibration light source and the spectral image of known spectral peaks associated with the light source. RainbowSTORM plots the line profile from the averaged background subtracted calibration image.

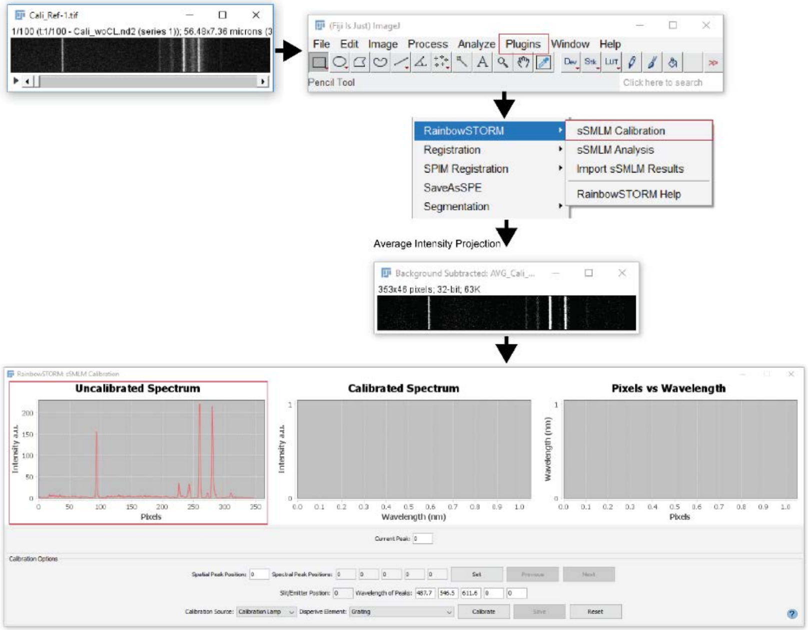

**Figure 1:** Calibration image processing workflow using RainbowSTORM

Next, the user can select the peaks to record the pixel position of the spatial and spectral peaks. The 'Current Peak' text field is updated by double clicking on the plot of the line profile or by manually inputting a value. The pixel position of the spatial or spectral peaks can be updated using the 'Set' button. The 'Next' and 'Previous' buttons can be used to toggle between the spectral peak position text fields. Additionally, the user can manually update the wavelength information of the peaks. By default, the wavelength information for 3 peaks at 487.7 nm, 546.5 nm and 611.6 nm from a calibration lamp are assigned. Next, the user specifies the calibration source and the dispersive element and presses the 'Calibrate' button to generate plots of the calibrated spectrum and the pixel to wavelength relationship. Pressing the 'Save' button saves the peak positions, the corresponding wavelengths, and the order of the calibration fitting polynomial as a CSV file. The blue question mark button launches RainbowSTORM help screen which provide the user with more information on the calibration screen.

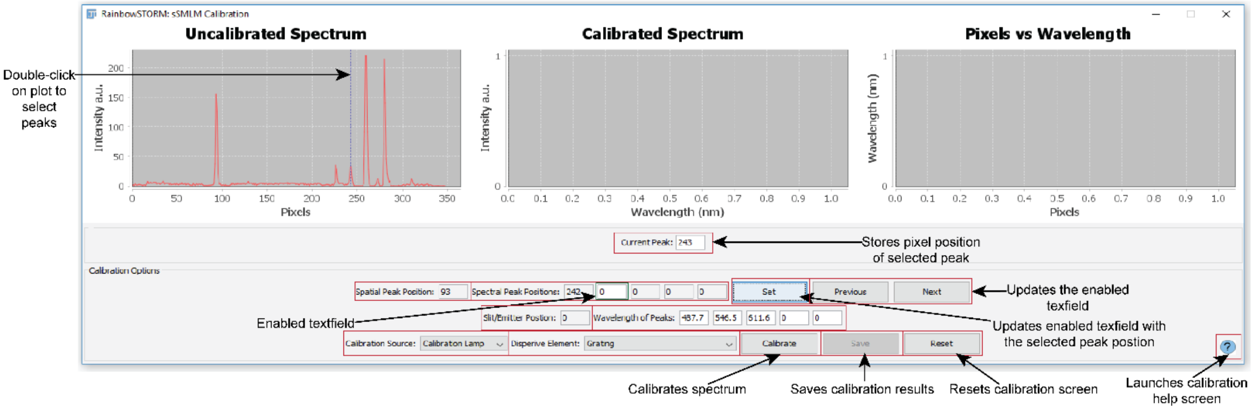

**Figure 2:** Calibration screen with the main features highlighted

##### 3.1.2 Fluorescent beads

Widefield images of fluorescent beads can be used in combination with a calibration light source to increase the number of points used for system calibration or to calibrate the system directly. Images of fluorescent beads feature their spatial locations and their corresponding spectra. Using the rectangular selection tool from the toolbar an individual bead and its corresponding spectra can be selected, and the associated frames duplicated. Optionally, for samples with uneven backgrounds the 'Subtract Background' option from the Process menu can be used as shown by the result in Step 2 of Figure 3. Using multicolor fluorescent beads (Figure 3), the emission maxima of the beads can be directly used to calibrate the system by following the same method used for calibration procedure using a calibration light source.

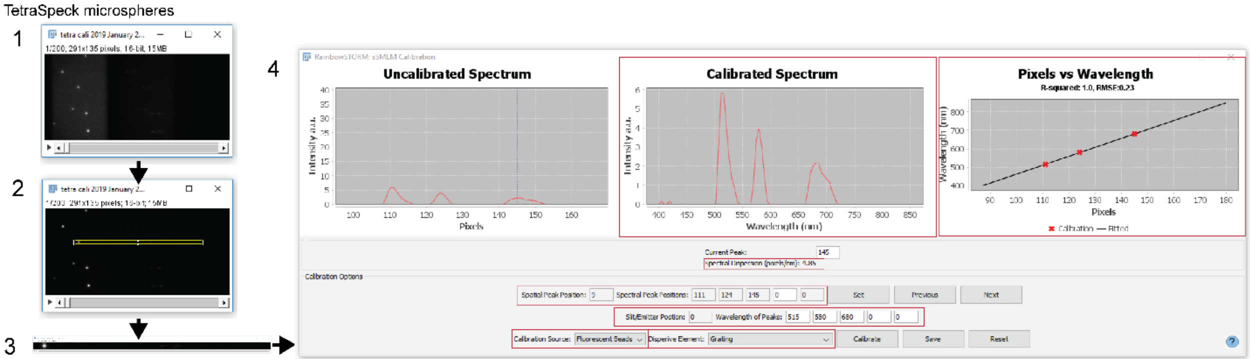

**Figure 3:** System calibration using multi-color fluorescent beads

Alternatively, using single-color beads (Figure 4) the pixel shift between the spatial and spectral pixel positions can be determined. This information can be manually entered to add additional points when calibrating the same system using a calibration light source (see section 3.2.2 for an example).

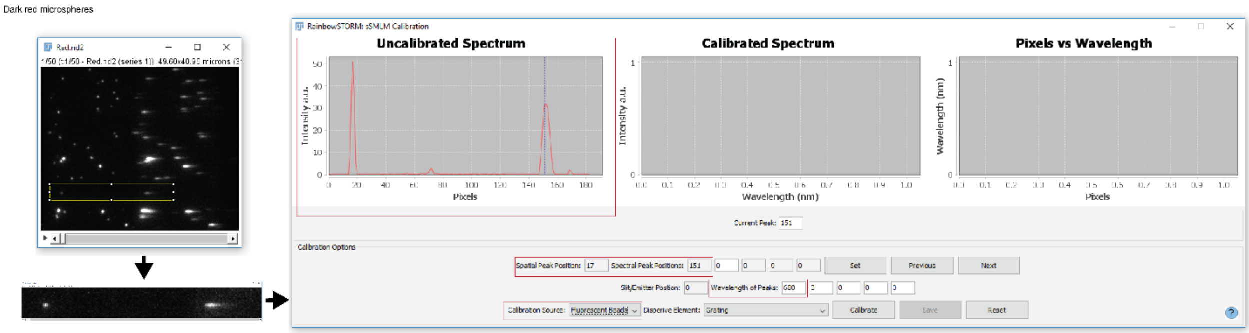

**Figure 4:** Calibration of a single-color fluorescent beads using RainbowSTORM

#### 3.2 Dispersive elements

sSMLM images can be recorded using a variety of optical designs. However, the key component in an sSMLM system is the dispersive element. Gratings and prisms can both be used to acquire spectroscopic information from single-molecule emitters. The choice of dispersive element impacts the fitting function used to calibrate the system shown in Figures 5 and 6. Since gratings disperse light linearly a first-order polynomial is used to relate the pixel positions to the wavelength information. In comparison, since prisms disperse light non-linearly a second-order or third-order polynomial can be used for calibration.

##### 3.2.1 Grating calibration

Using a grating as the dispersive element, the pixel positions and corresponding wavelengths are fit to a first-order polynomial. RainbowSTORM requires at least three data points for accurate fitting. The goodness of fit ($R^2$) and root mean square error (RMSE) for the points are shown in the pixel versus wavelength plot.

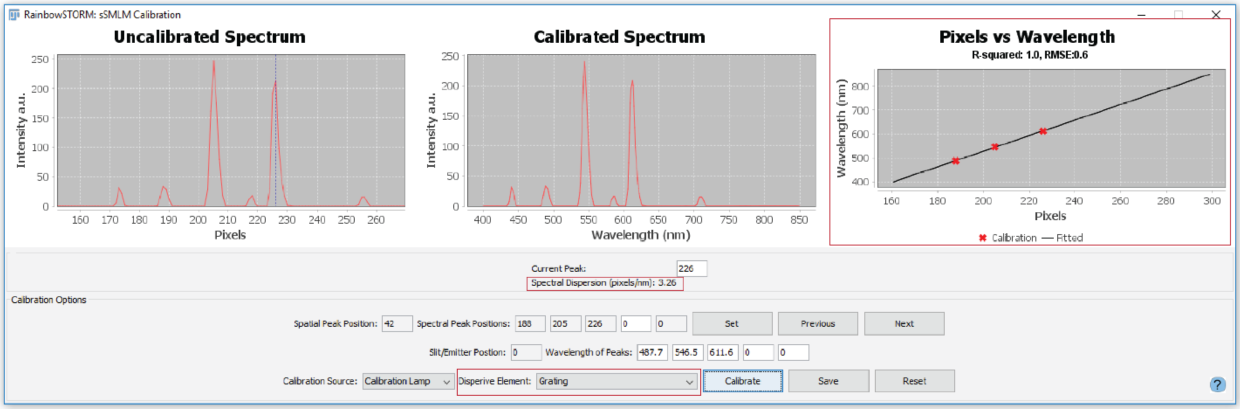

**Figure 5:** Calibration results of a grating-based system using RainbowSTORM

Additionally, differences in the pixel positions along the field of view, which can indicate potential alignment imperfections, are automatically detected and reported when using a calibration light source.

##### 3.2.2 Prism calibration

Using a prism as the dispersive element, the pixel positions and corresponding wavelengths are fit to a second-order or third-order polynomial. RainbowSTORM requires at least four data points for accurate fitting; however, five are recommended for third-order polynomial fitting. In the example below, three laser lines are used as the calibration light source in combination with spectroscopic information from single emitter fluorescent beads. The goodness of fit ($R^2$) and root mean square error (RMSE) for the points are shown in the pixel versus wavelength plot.

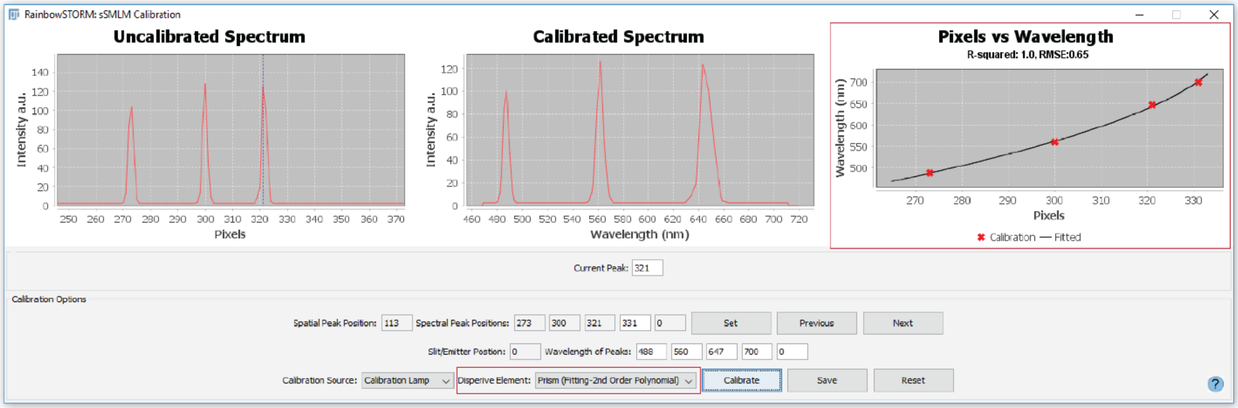

**Figure 6:** Calibration results of a prism-based system using RainbowSTORM

After calibrating the sSMLM system, experimental images can be loaded into ImageJ. RainbowSTORM's sSMLM analysis screen (Figures 7 and 8) can be launched from the RainbowSTORM Menu. Using this screen sSMLM images are separated into spatial and spectral images. While the spatial images are analyzed using the existing SMLM analysis plugin (ThunderSTORM), all spectroscopic analysis is performed using RainbowSTORM. After spatial analysis, RainbowSTORM removes background signals from spectral images and relates each single-molecule localization to its corresponding spectral image. Users can also define the parameters to visualize sSMLM images. RainbowSTORM then identifies the spectral signature for each localization from the spectral images using the spatial coordinates of each localization and the spectral calibration information. Additionally, RainbowSTORM calculates the weighted spectral mean (spectral centroid), photon count in the spectral image, background photon count in the spectral image, background photon count per pixel in the spectral image, and spectral precision of each localization. Using RainbowSTORM's 'Process sSMLM Data' module, the wavelength range for spectral analysis can also be specified. Users can also elect to automatically reject localizations with overlapping spectral images. Additionally, using the advanced settings the width of the spectrum which by default is estimated by fitting the line spectrum of each spectral image to a gaussian can be defined by the user. RainbowSTORM's advanced settings also allows users to define the number of pixels along the y-axis of the spectral image (localization pixel shift) can be specified. Differences between the y position of the spectral image and the spatial image of each localization can also be compensated using RainbowSTORM. Before running the sSMLM analysis, the spectral images and extracted line spectra can be previewed by pressing the 'Preview' button. This allows users to adjust the background subtraction or any other spectroscopic analysis parameter before processing all the data. Users can also reset all the parameters in the processing menu to the default parameters by pressing the 'Reset' button. The help screen for the analysis screen can be accessed by pressing the blue question mark button. Once users are satisfied with the settings, they can select the 'Run Analysis' button to process the spectroscopic data.

### 4. sSMLM Analysis

The examples below show how sSMLM data can be processed using RainbowSTORM.

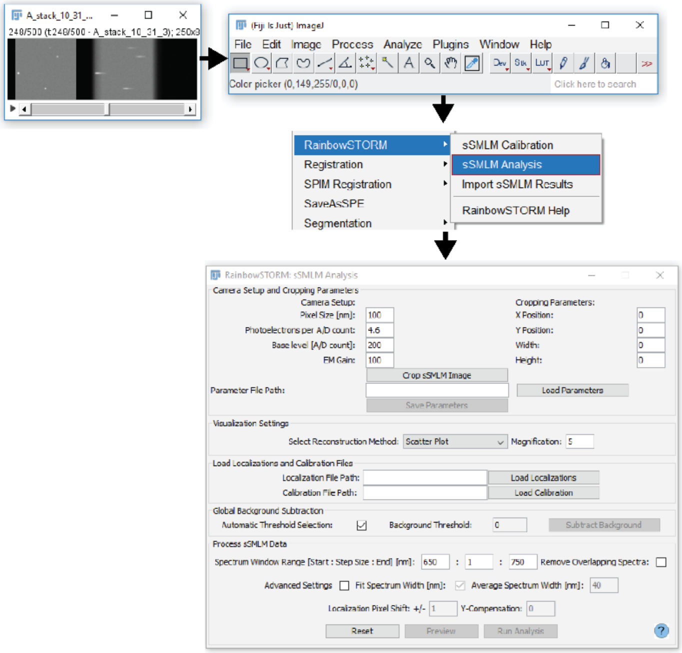

**Figure 7:** Launching RainbowSTORM's analysis screen

#### 4.1 Process spatial information

sSMLM analysis requires the localization of single molecules and the analysis of their corresponding spectral signature. This is done using the sSMLM analysis screen shown below.

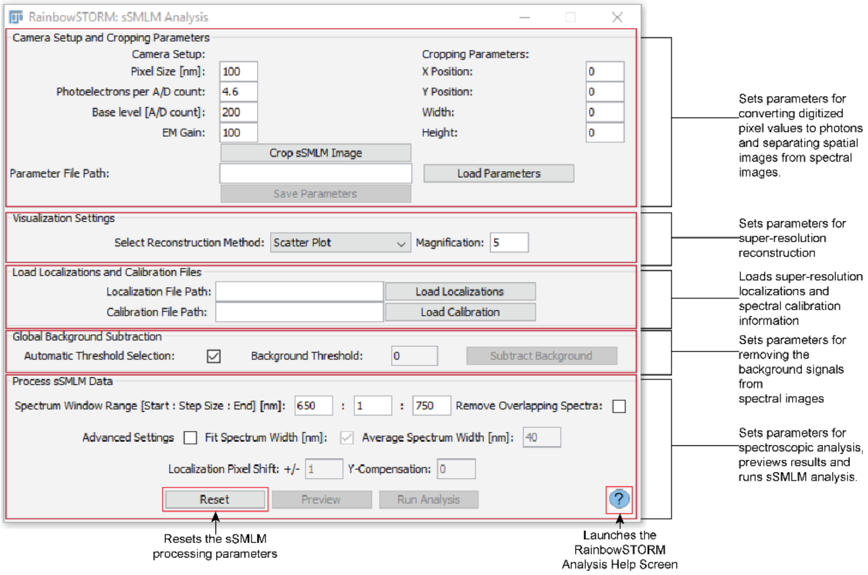

**Figure 8:** RainbowSTORM's sSMLM analysis screen with the main analysis modules highlighted

##### 4.1.1 Set camera parameters

The camera pixel size in nanometers, conversion factor between photons and digital units, base level offset of the camera in digital units, and electron multiplier gain of the camera must be set for conversion from digital units to photons. The camera setup options are shown in Figure 9.

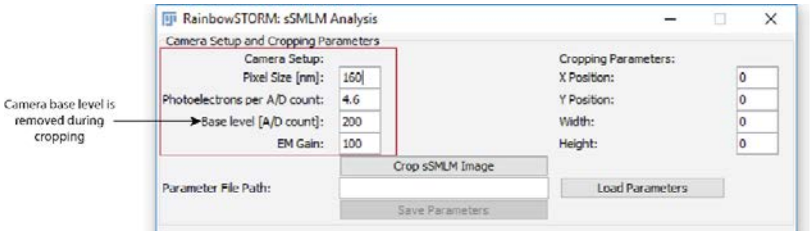

**Figure 9:** RainbowSTORM's camera setup and cropping module with the camera setup parameters

##### 4.1.2 Crop image and remove base level offset

To prepare the recorded sSMLM data for spatial analysis, the images are split into spatial and spectral images using RainbowSTORM's cropping tool as shown in Figure 10. The cropping information can be updated using ImageJ to select a rectangular region of interest (ROI) and pressing the 'Crop sSMLM Image' button. This updates RainbowSTORM's cropping information (initial x and y pixels and the rectangle width and height) and generates the spatial images with the selected dimensions. The ROI information can also be manually updated. Next, RainbowSTORM generates the spectral image with the same height as the spatial image but the width is determined by the end of the cropped spatial image and the width of the original sSMLM image. The camera base level is also removed from the spectral images.

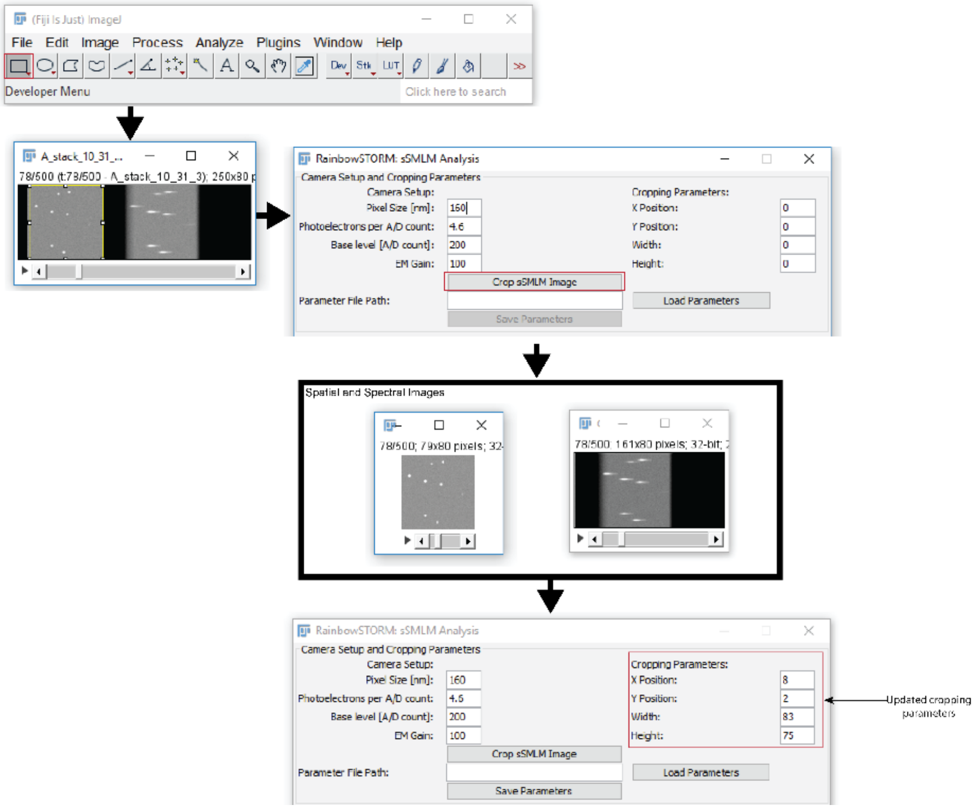

**Figure 10:** Workflow for separating sSMLM images into spatial and spectral images using the RainbowSTORM cropping module.

The 'Save Parameters' button saves the current camera information and the cropping parameters as a CSV file for future use. Previously, saved parameters can be loaded by pressing the 'Load Parameters' button and selecting the file or by inputting the file location and name and then pressing the 'Load Parameters' button.

##### 4.1.3 Process SMLM Data using ThunderSTORM

The cropped spatial image can then be analyzed using ThunderSTORM (shown in Figure 11), a single molecule localization tool. First the ThuderSTORM camera settings should be updated using the RainbowSTORM camera settings. Next, the ThunderSTORM parameters can be updated as outlined in ThunderSTORM's manual and processed.

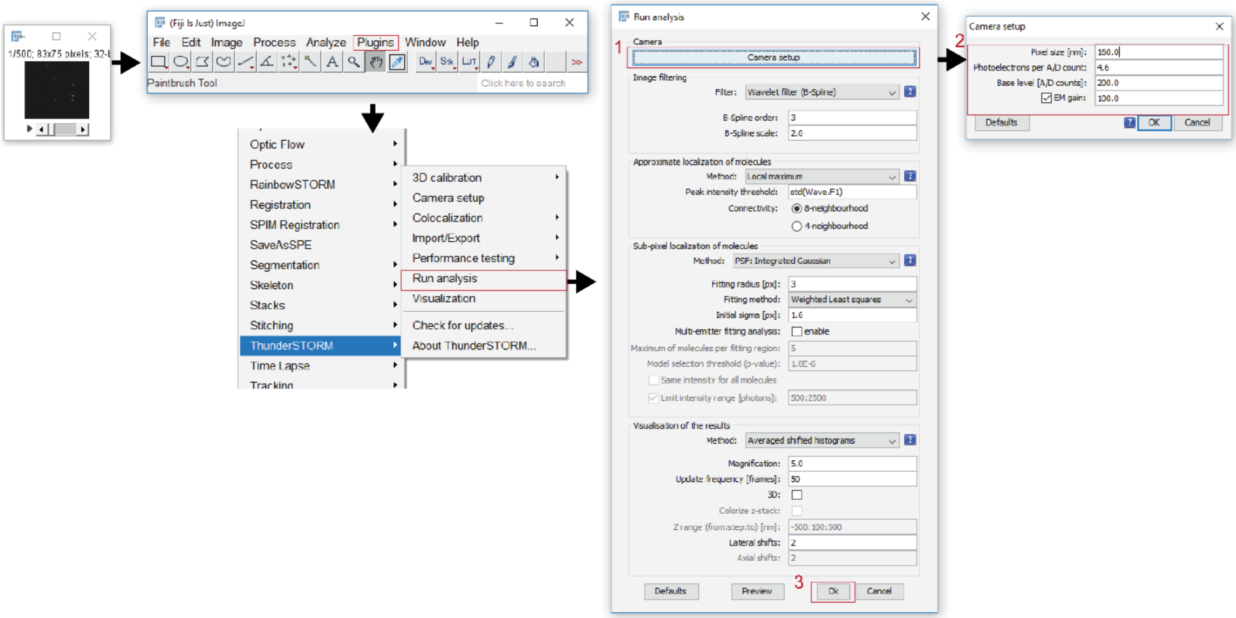

**Figure 11:** Workflow for processing spatial images using ThunderSTORM

The ThunderSTORM coordinates should be in nanometers, if not click on the header of the results table (shown in Figure 12) to set the units. Users can then select the 'Export' button. Finally, the resulting SMLM results can be saved as a CSV.

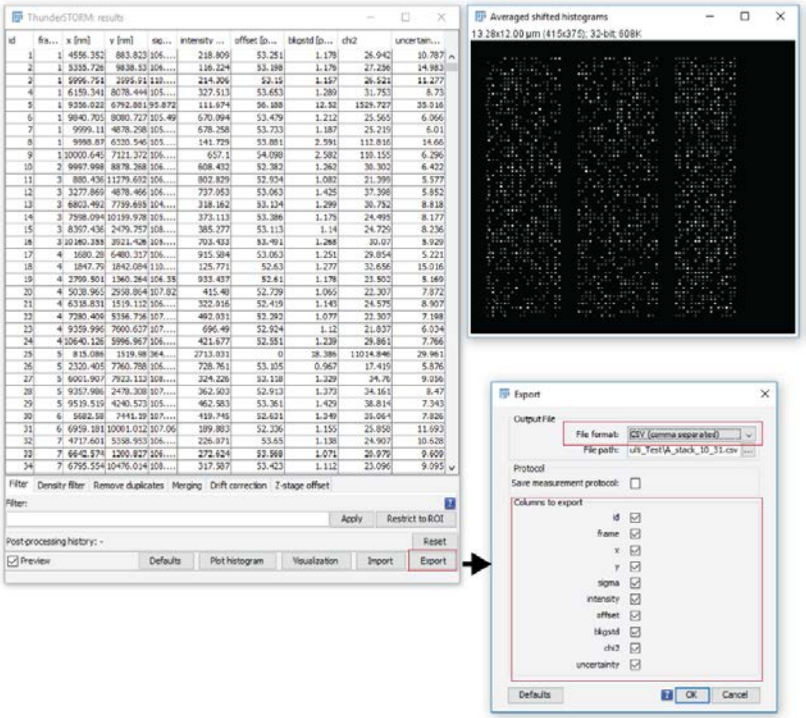

**Figure 12:** Workflow for exporting ThunderSTORM results

Additionally, the SMLM results can be drift corrected using ThunderSTORM (shown in Figure 13) and saved as a JSON file for later use by RainbowSTORM. Note that RainbowSTORM requires localizations without drift-correction. The localizations can be drift-corrected on the visualization screen after spectroscopic analysis.

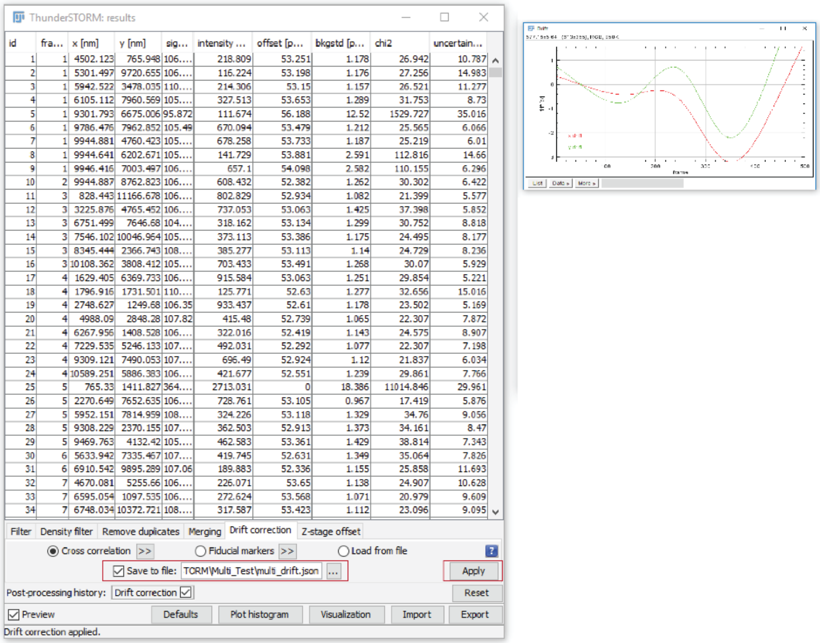

**Figure 13:** Drift-correction of ThunderSTORM results

Three-dimensional (3D) sSMLM images (Figure 14) can be acquired by placing a cylindrical lens in the spatial imaging path. Prior to acquiring the sSMLM images the spatial channel can be calibrated using fluorescent beads and processed using ThunderSTORM (Ovesny, et al., 2014). Using RainbowSTORM the spatial and spectral images can be separated. The 3D spatial images can then be processed using the ThunderSTORM parameters for 3D astigmatism analysis (Ovesny, et al., 2014). This process will add the following spatial fields to the analysis: z, sigma1, and sigma2.

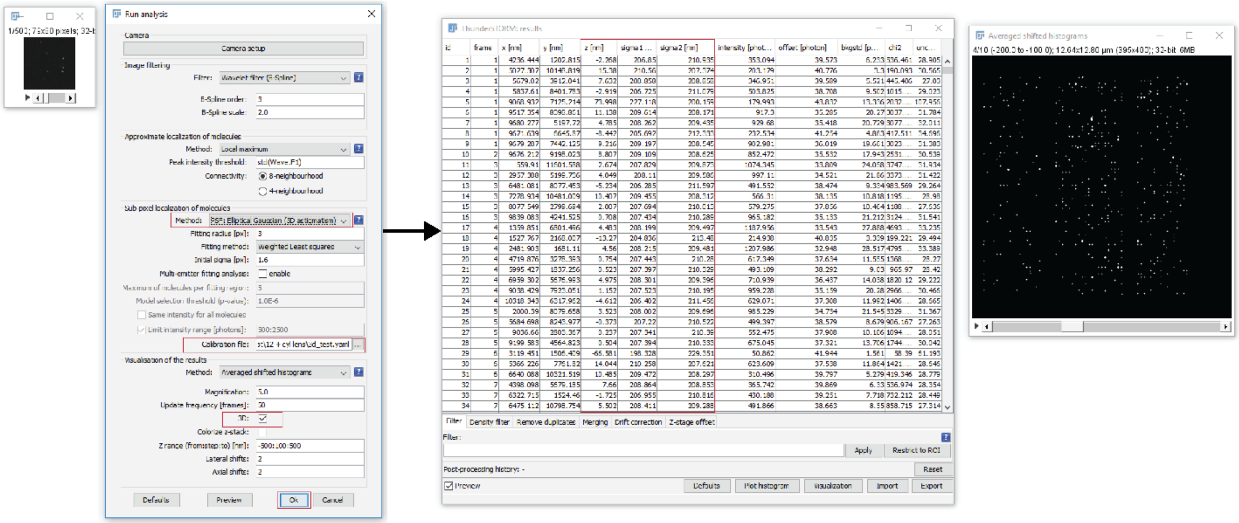

**Figure 14:** ThunderSTORM 3D astigmatism processing workflow and results

After saving the localization information, ThunderSTORM can be closed, and all other processing handled using RainbowSTORM.

#### 4.2 Set Visualization settings

The visualization settings module (Figure 15) allows the reconstruction method and magnification of the reconstructed image to be set. RainbowSTORM renders super-resolution images using scatterplots or as averaged gaussian plots. Scatterplots show the location of each emission event as equally weighted points. Averaged gaussian plots show the location of each emission event as a 2D gaussian whose radius reflects the localization uncertainty of the emission event.

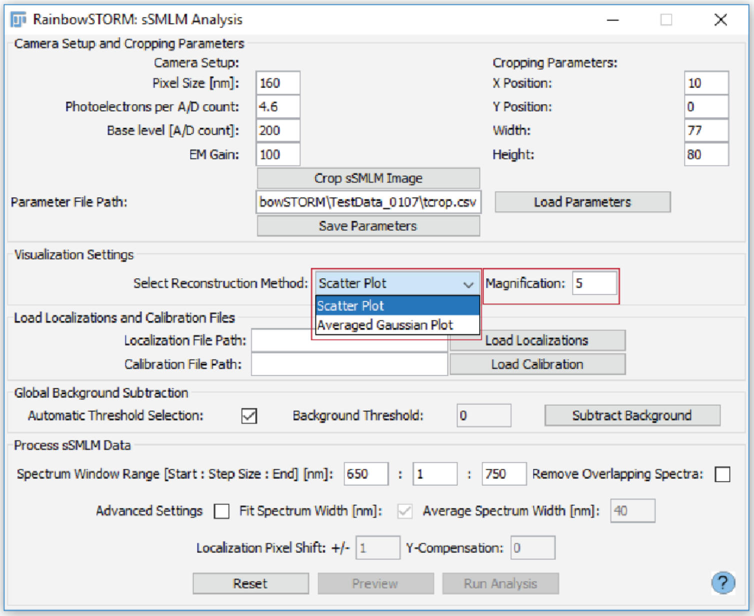

**Figure 15:** RainbowSTORM's visualization settings

#### 4.3 Load SMLM and calibration data

Next the SMLM results and calibration CSV files must be loaded into RainbowSTORM as shown in Figure 16. After loading the SMLM results, the SMLM image will be reconstructed based on RainbowSTORM's visualization settings.

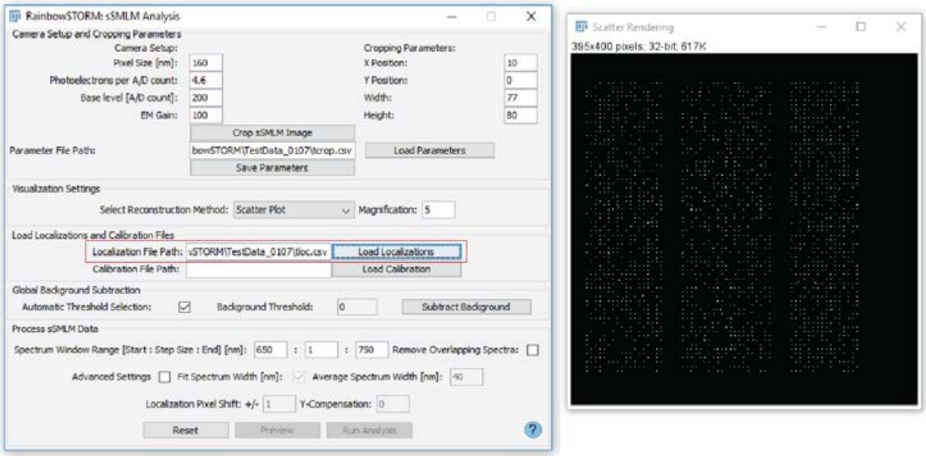

**Figure 16:** Workflow for loading saved ThunderSTORM localizations using the sSMLM analysis screen

The pixel versus wavelength plot is output when the calibration file is successfully loaded as shown in Figure 17.

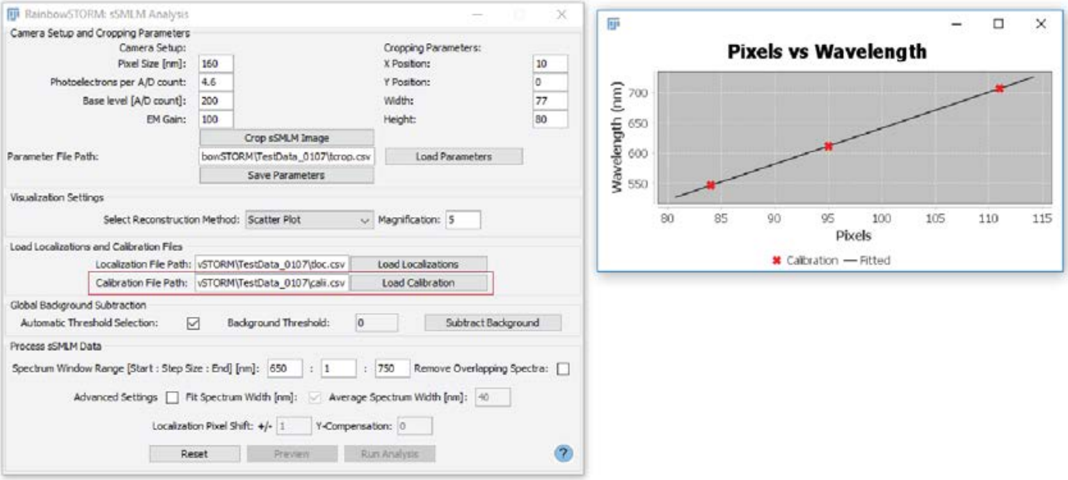

**Figure 17:** Workflow for loading saved calibration information using the sSMLM analysis screen

#### 4.4 Background subtraction

The background signals from the spectral images can be removed using global background subtraction. The threshold for global background subtraction can be automatically generated or user defined. To determine the global background, each row of the image is selected and the average pixel value over the frames of the stack is recorded for each pixel and saved. The thresholds are used to reject pixels, which contain signals from the sample. The remaining signals are averaged and used to create a background image which is subtracted from each of the spectral images. The preview panel can be used to view the spectral image for individual localization events and their spectral profile. If the background has not been effectively removed, the threshold can be manually adjusted.

##### 4.4.1 Automatic thresholding

By default, automatic thresholding (Figure 18) is selected; however, this can be deselected by unchecking the checkbox in the background subtraction module. Pressing the ‘Subtract background’ button results in automatic generation a threshold for each row in the spectral image. To automatically set the threshold the average digitized pixel values for all the frames of the image for the selected row are analyzed. From these averaged pixel values the minimum pixel and the standard deviation of the pixels are calculated. The threshold for that row is the sum of the minimum pixel value and four times the standard deviation. The threshold is used to assess each pixel across the frames of the spectral image. Pixels which are less than the threshold are saved, and the average of the selected pixels are used to create the global background image. The background image is then subtracted from each frame of the spectral images. Once the process is completed, a background image and a stack of background-subtracted spectral images are displayed.

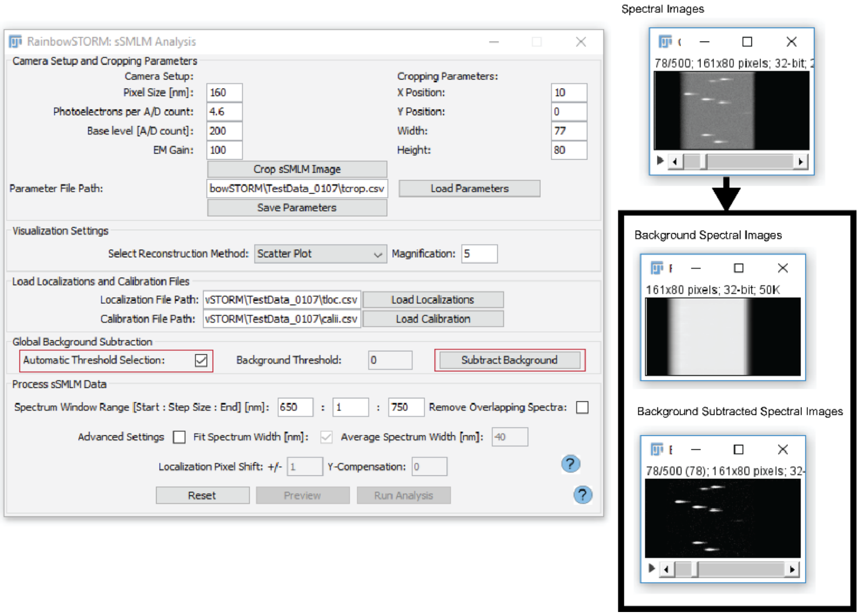

**Figure 18:** Workflow for background subtraction using RainbowSTORM's automatic background threshold

#### 4.4.2 Manual thresholding

To manually select a threshold the automatic thresholding checkbox must be unselected. Once this is done, the global background threshold text field will be enabled. Digitized signal levels from the background of the spectral images can be determined by selecting background pixels in the spectral image. These values can be used to set an appropriate threshold for background subtraction as shown in Figure 19. The 'Subtract Background' button can be pressed to generate a background image and a stack of background-subtracted spectral images.

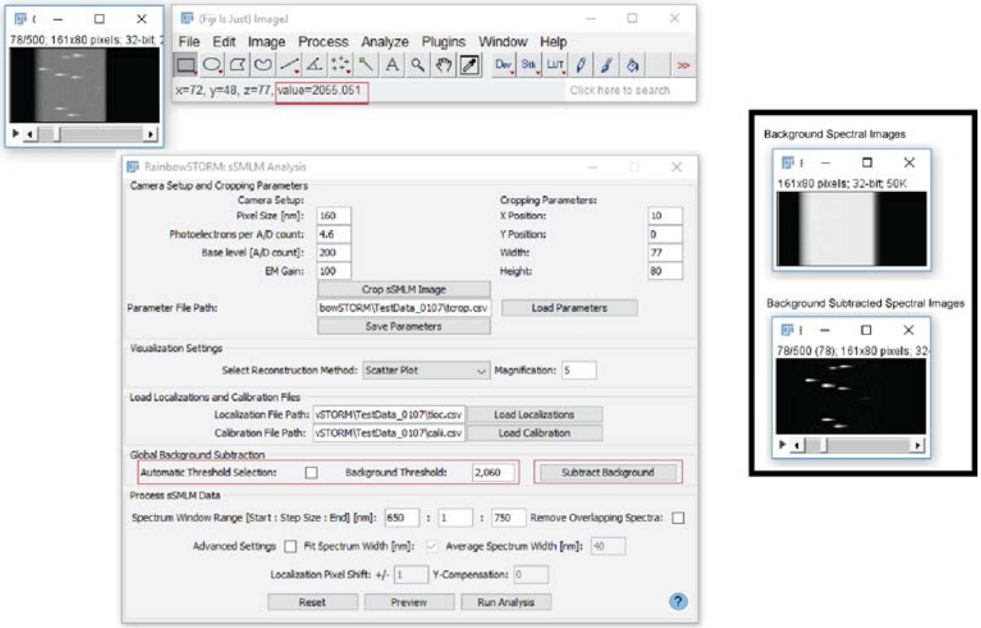

**Figure 19:** Workflow for background subtraction using a user-defined background threshold

#### 4.5 Process spectroscopic data

To process the spectra for each localization event the spectral window range is updated based on the fluorescence emission of the dyes used in the sSMLM experiment. The preview button can be selected to observe the results for each of the emission events. The main features for processing sSMLM data are highlighted in Figure 20.

##### 4.5.1 Set processing parameters

The main parameters for spectroscopic analysis are the start, end and step size for the spectrum window range. Additionally, users can opt to remove localizations which overlap in space. The spectrum window range can be set based on the fluorescence emission of the single molecule dyes used in the experiment. We recommend a spectral window bandwidth from 100 nm to 150 nm. More advanced users can select the ‘Advanced Settings’ checkbox to enable modification of the spectra width in nanometers, localization pixel shift and y-compensation fields. The spectrum width can be calculated by fitting the line spectrum to a gaussian, alternatively the user can manually input the average spectrum width for the dyes being used. By default, the automatic spectrum width option is enabled, to manually set the spectrum width, uncheck the ‘Fit Spectrum Width’ checkbox. The localization pixel shift controls the number of pixels required to cover the spectral image along the y-axis. The localization pixel shift also captures the number of pixels to cover the point spread function of each localization along the x and y axes. By default, this is set to ±1 for a total of 3 pixels; however, this can range from ±1 to ±3 for a total of 3 to 7 pixels for each point spread function. The y-compensation field can be used to adjust the pixel position of the spectral images along the y axis. By default, the y-compensation is set to 0 but can range from -5 to 5. All parameters on the processing panel can be reset to the defaults by selecting the reset button.

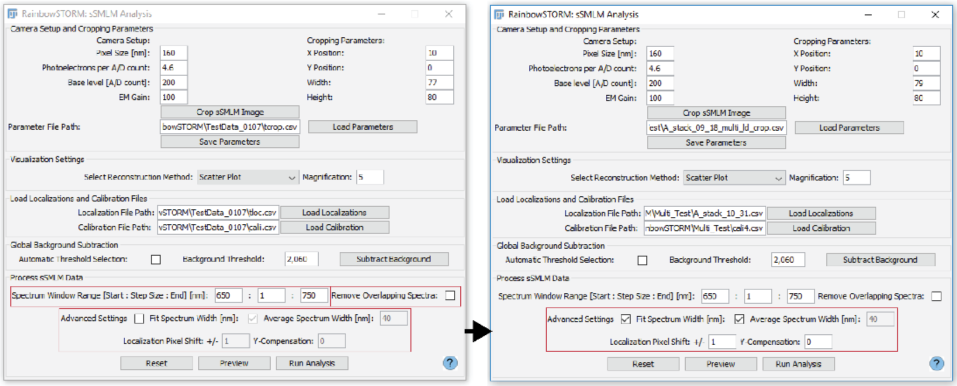

**Figure 20:** RainbowSTORM’s sSMLM processing module with the main features highlighted

##### 4.5.2 Preview sSMLM data

The ‘Preview’ button allows for the results of the current settings to be previewed for up to 100,000 localization events. The ‘Next Event’ and ‘Previous Event’ buttons switch between localization events. The ‘Update Event’ button reprocesses the current localization event using updated background subtraction and spectroscopic processing parameters. The preview window (Figure 21) also shows the percentage of events, which would be removed by the automatic overlapping spectra removal described in the following section.

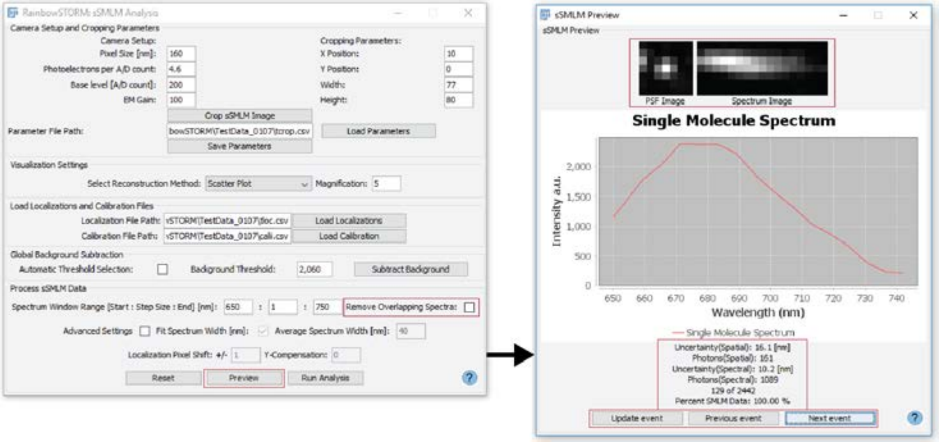

**Figure 21:** RainbowSTORM's preview screen

#### 4.5.3 Remove overlapping spectra

For accurate spectroscopic analysis, localizations with spectral images which overlap in space should be removed. This can be done by automatically rejecting the overlapping localizations by selecting the 'Remove Overlapping Spectra' checkbox. The results can be viewed using the preview screen as shown in Figure 22. Alternatively, these localizations can be filtered out by using post-processing filters (e.g. point spread function size and spectrum width).

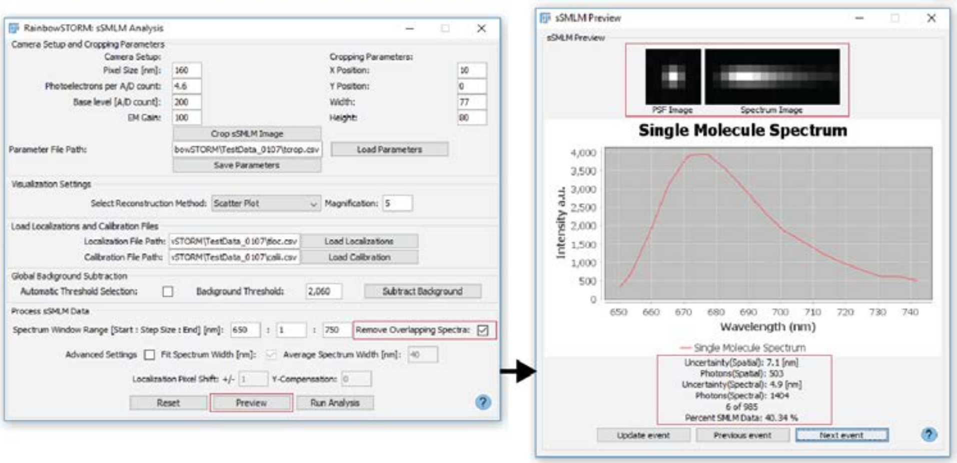

**Figure 22:** RainbowSTORM's preview screen after removing localizations with overlapping spectra

#### 4.5.4 Analyze sSMLM images

Once the spectroscopic analysis parameters are set, users can press the 'Run Analysis' button to process the spectra for all localized emission events. During processing, the ImageJ status and progress bars are updated. After analysis, the Pseudo-colored sSMLM super-resolution image reconstructions are rendered and the spectroscopic results for all emission events will be shown on the sSMLM Visualization screen. The workflow and results of this sSMLM analysis are shown in Figure 23.

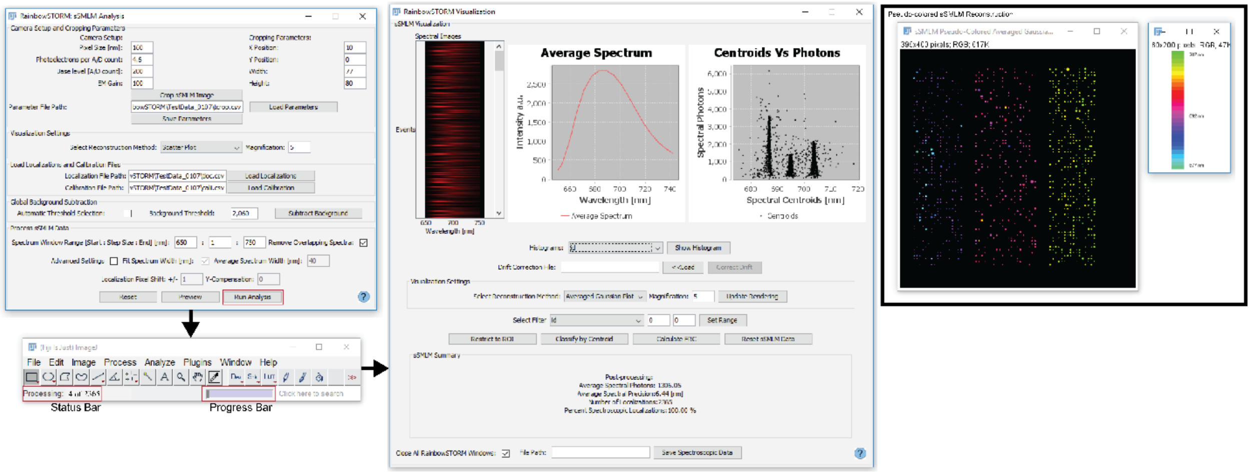

**Figure 23:** Workflow for processing sSMLM images and results of spectroscopic analysis

### 5. sSMLM visualization and post-processing

The sSMLM visualization screen (Figure 24) shows images of the emission spectra, a line plot of the averaged spectrum, and a scatter plot of the calculated spectral centroids versus the total photon counts of the spectral signatures. The spatial and spectral fields (Table 1) can be displayed by selecting the field from the combination box and pressing the ‘Show Histogram’ button. ThunderSTORM drift-correction files can be loaded and applied by selecting the ‘Correct Drift’ button. The reconstruction method and magnification can be updated in the visualization settings module. Using the corresponding histograms, the fields can be filtered by specifying a range, in which the values for that field should fall. Table 1 includes descriptions of all RainbowSTORM fields. Unless, otherwise specified the acceptable range for filtering is limited to positive integers.

Additionally, regions of interest (ROI) in the images can be selected using the rectangular selection tool and the updated reconstructions rendered by selecting the Restrict to ROI’ button. All filters can be removed, and the original results reloaded by selecting the ‘Reset sSMLM Data’ button. The sSMLM summary panel shows the average photon count and spectral precision, the number of localizations as well as the percentages of the total sSMLM and the total SMLM results included in the reconstruction. Post-processing steps are also tracked using the summary panel. The ‘Classification’ button launches the classification screen which can be used to generate multicolor super-resolution images based on localizations within one or more specified centroid windows. Clicking the ‘Save’ saves the results into a CSV file. Finally, users who wish to keep RainbowSTORM screens open after closing the visualization screen can uncheck the ‘Close All RainbowSTORM Windows’ checkbox. The help screen for the visualization screen can be accessed by pressing the blue question mark button.

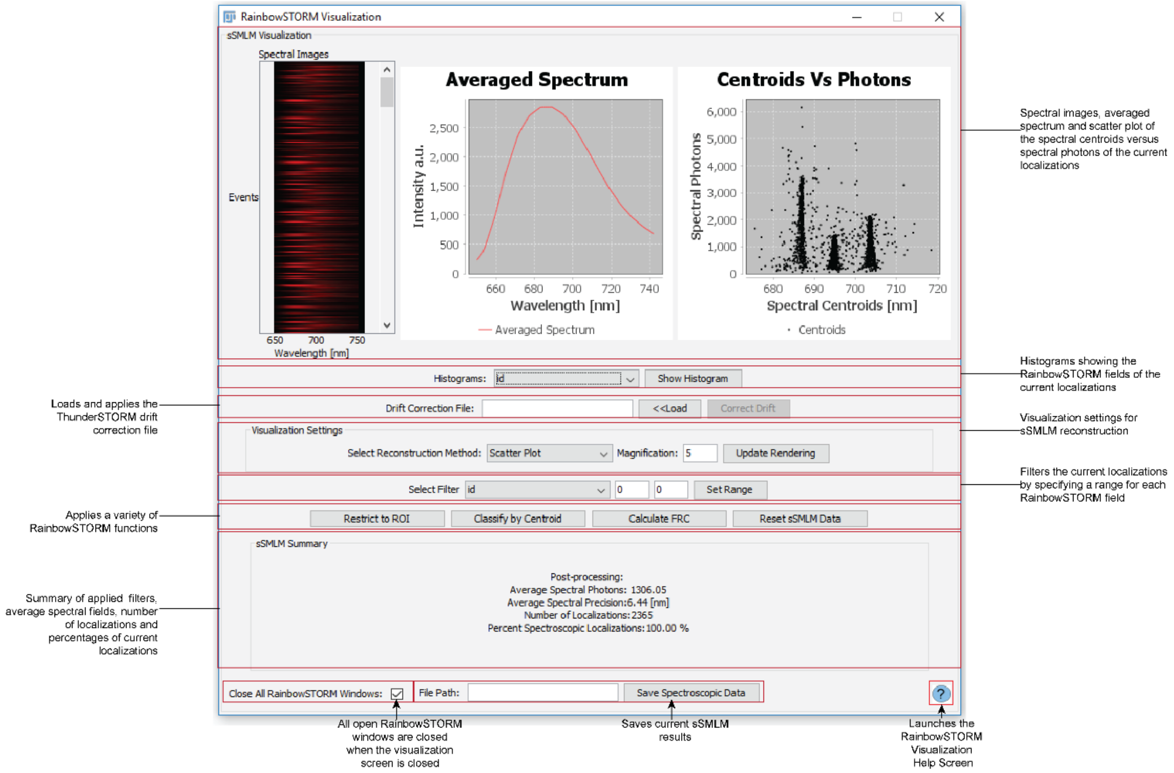

**Figure 24:** RainbowSTORM's visualization screen with the main features highlighted

Table 1: Descriptions of each RainbowSTORM field.

| **Field** | **Description** |
|---|---|
| id | Index of the current fitted blinking events. |
| frame | The frame of each blinking event. Set using ThunderSTORM. |
| x | The subpixel x position of each blinking event. Used to reconstruct the super-resolution images. Values are confined within the bounds of the original image or selected region of interest. Estimated using ThunderSTORM. |
| y | The subpixel y position of each blinking event. Used to reconstruct the super-resolution images. Values are confined within the bounds of the original image or selected region of interest. Estimated using ThunderSTORM. |
| z | The subpixel z position of each blinking event estimated using the astigmatism processing method using ThunderSTORM. Used to reconstruct the 3D super-resolution images. Accepted range: -2000 nm to 2000 nm. |
| spatial photons | Total photons of the fitted blinking event in the spatial domain. Used to estimate the localization uncertainty. Estimated using ThunderSTORM. |
| spatial sigma | The standard deviation of the fitted blinking event in the spatial domain. Used to estimate the localization uncertainty and point spread function width along the focal plane. Estimated using ThunderSTORM. |
| spatial sigma1 | The standard deviation of the fitted blinking event in the spatial domain along the lateral axis for 3D sSMLM images. Used to estimate the localization uncertainty along the lateral axis and point spread function width along the focal plane. Estimated using ThunderSTORM. |
| spatial sigma2 | The standard deviation of the fitted blinking event in the spatial domain along the axial axis for 3D sSMLM images. Used to estimate the point spread function width along the axial axis. Estimated using ThunderSTORM. |

| | |
|---|---|
| localization uncertainty | The estimated uncertainty in the lateral position of each fitted blinking event. Combined with sample labeling, the localization uncertainty is used to estimate the resolution of the reconstructed image. Estimated using ThunderSTORM. |
| spectral centroid | The spectral centroid or spectral mean of each blinking event. The centroid can be used to infer information related to the identity, local environment and conformational state of the source (dye or protein) of the blinking event. Values are used to color-code pseudo-colored super-resolution images. Estimated using RainbowSTORM. Accepted centroids range from 380 nm to 870 nm with a minimum step size of 1. |
| spectral photons | The total photons of the spectrum of each blinking event. Used to estimate the spectral uncertainty. Estimated using RainbowSTORM. |
| spectral bk photons | The total background photons of the spectrum of each blinking event. Estimated using RainbowSTORM. |
| spectral bk photons/pixel | The total background photons for each pixel of the image of the spectrum of each blinking event. Used to estimate the spectral uncertainty. Estimated using RainbowSTORM. |
| spectral sigma | The standard deviation of the spectrum width. This value is estimated from fitting the spectrum to a Gaussian or using the average spectrum width for a given source. Used to estimate the spectral uncertainty. Estimated using RainbowSTORM. |
| spectral uncertainty | The estimated uncertainty in the spectral centroid of each blinking event. This can be used to exclude events whose spectral centroid cannot reliably be identified. Estimated using RainbowSTORM. |

#### 5.1 Pseudo-colored image reconstructions

Upon launching the visualization screen, a Pseudo-colored sSMLM super-resolution reconstruction (Figure 25) and its corresponding colorbar are displayed. The colors in the image are selected based on the calculated centroid. The range of the colorbar is set by the centroid distribution. Updating the sSMLM results also updates the Pseudo-colored reconstructions accordingly.

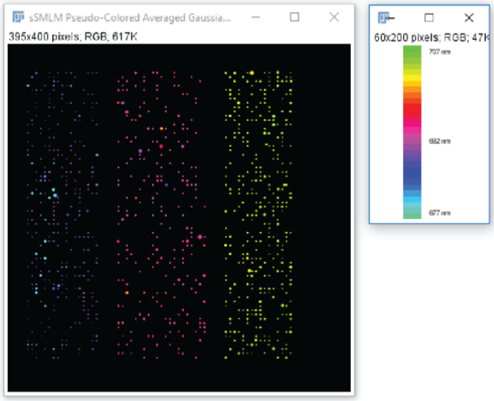

**Figure 25:** Pseudo-colored super-resolution image reconstructions and corresponding colorbar

#### 5.2 sSMLM filtering

Post-processing filters are commonly used in both SMLM and sSMLM. RainbowSTORM's visualization screen allows users to select specific segments of the data or remove outliers by using both the spatial and spectral fields to apply filters. Additionally, ROIs in the super-resolution reconstruction can be selected and analyzed independently. All filters can be removed and the original sSMLM data can be reloaded by clicking the 'Reset sSMLM Data' button.

#### 5.21 sSMLM histograms

Prior to applying post-processing filters, the distribution of the spatial and spectral fields can be visualized by displaying their histograms as shown in Figure 26. This is accomplished by selecting individual fields from the histogram combination box. Pressing the 'Show Histogram' button launches an ImageJ histogram window showing the distribution of the selected field. Using this information, users can set appropriate filters to select specific sections of the data or remove any outliers. The example below demonstrates this process.

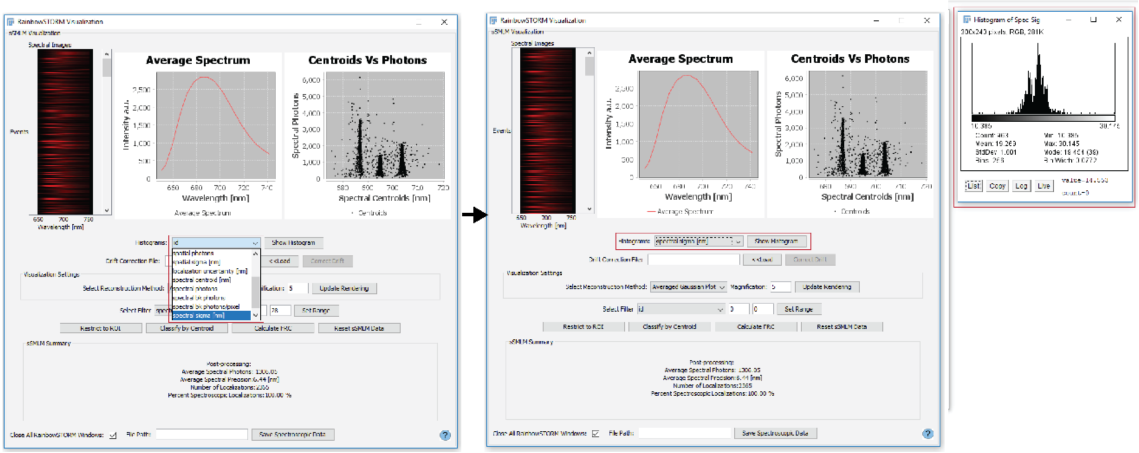

**Figure 26:** Workflow to show the histograms of a selected sSMLM field

#### 5.22 Filtering data using sSMLM fields

Once users understand the distribution of the field, which will be used for filtering the sSMLM data, a range for the localizations of interest can be specified using the filter options on the visualization screen (Figure 27). This is done by first selecting the field from the filter combination box, then the start and end points for the range are specified using the corresponding text fields. RainbowSTORM detects any invalid start and end points and informs the user of the acceptable inputs. Next the 'Apply Filter' button is pressed, resulting in the localizations displayed in the visualization panel being restricted based on the filter parameters. The Pseudo-colored image reconstruction will be updated using the restricted localizations. The summary results will also be updated to show the filter which has been applied as well as new statistics for the updated localizations.

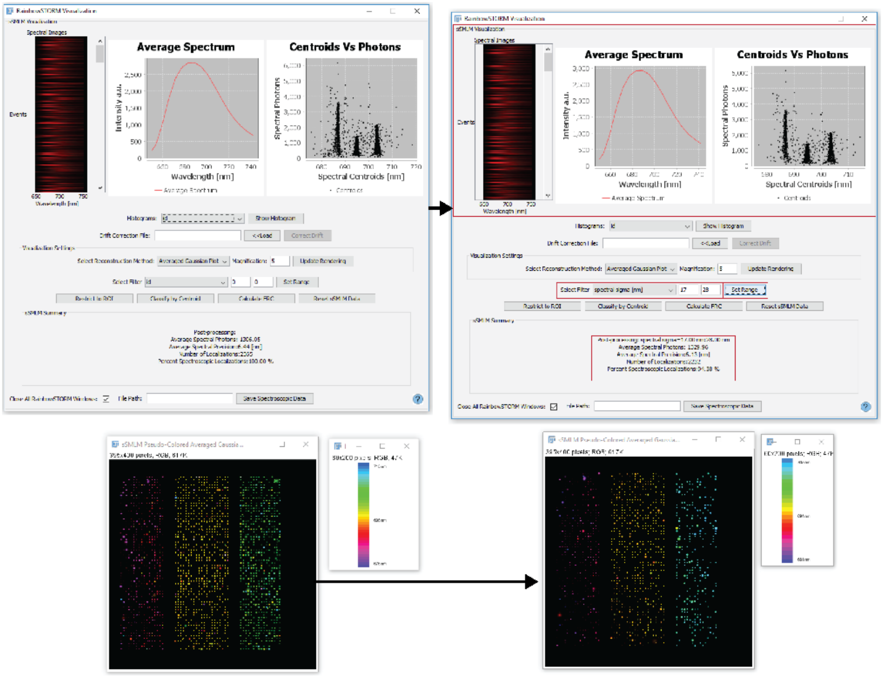

**Figure 27:** Workflow showing RainhowSTORM filtering by a selected field as well as the updated visualization screen and sSMLM super-resolution reconstructions

##### 5.23 Restrict to ROI

RainbowSTORM also allows user to restrict analysis to only specific ROIs in the sSMLM super-resolution reconstruction as shown in Figure 28. This can be done using ImageJ's rectangular selection tool to select the ROI then pressing the 'Restrict to ROI' button. This will result in an updated Pseudo-colored image being rendered, and the localizations displayed on the visualization screen will be updated. In addition, the post-processing tracker in the summary panel will be updated to show the application of this filter.

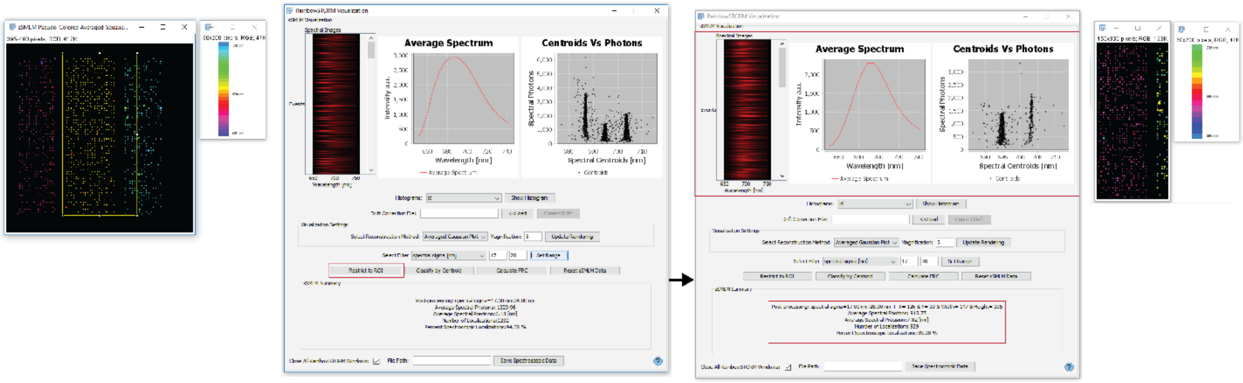

**Figure 28:** Workflow for filtering localizations by selecting a region-of-interest in the sSMLM super-resolution image with the updated visualization screen and Pseudo-colored super-resolution reconstruction

#### 5.2 Classification

Clicking the classification button launches the classification screen (Figure 29). Up to six (6) color spectral windows can be assigned to predefined color channels based on the centroid distribution. The 'Show Channel' checkbox can be selected to show each single-color channel. Selecting the 'Classify by Centroid' button launches a multicolor image of the selected data and a channel summary (a scatter plot of the centroids versus photons and a histogram of the centroid distribution) with the channels highlighted. The channel summaries are optional and can be deselected using the ‘Show Chanel Summary’ checkbox. The help screen for the classification screen can be accessed by pressing the blue question mark button.

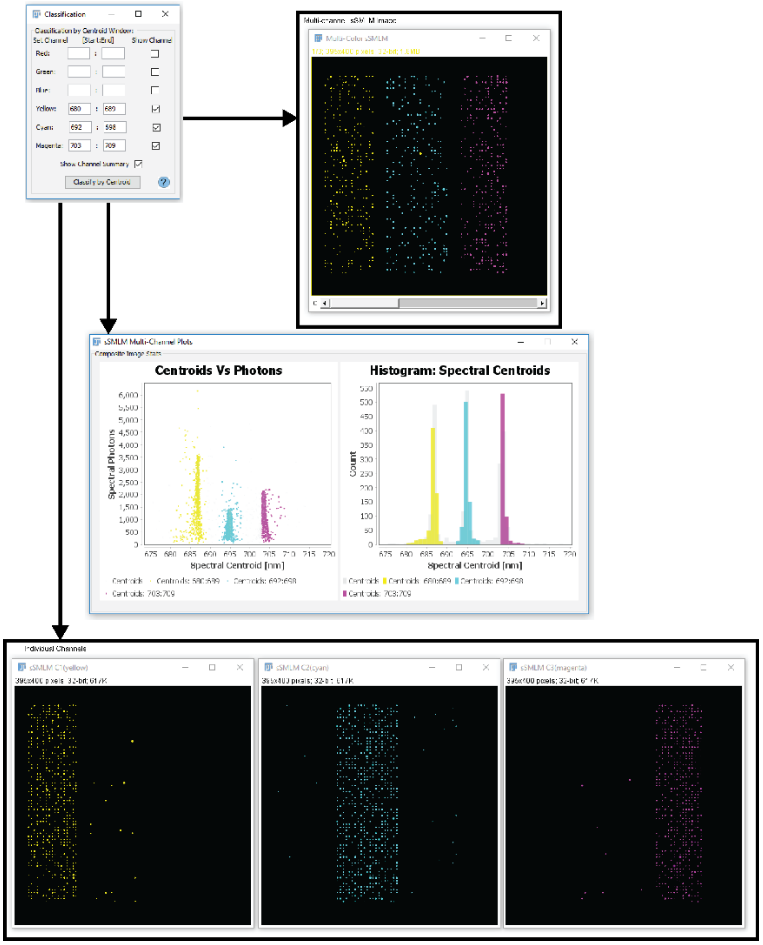

**Figure 29:** Workflow for classifying localizations by their spectral centroids as well as multicolor super-resolution reconstructions and plots of the results.

#### 5.3 Three-dimensional (3D) sSMLM rendering

A stack of 3D sSMLM images (Figure 30) can be generated using RainbowSTORM by selecting the ‘Display 3D sSMLM’ button on the visualization screen. The average spatial sigma2 is used to set the step size along the axial position. This button is only available if RainbowSTORM detects the z position field when ThunderSTORM localizations are loaded.

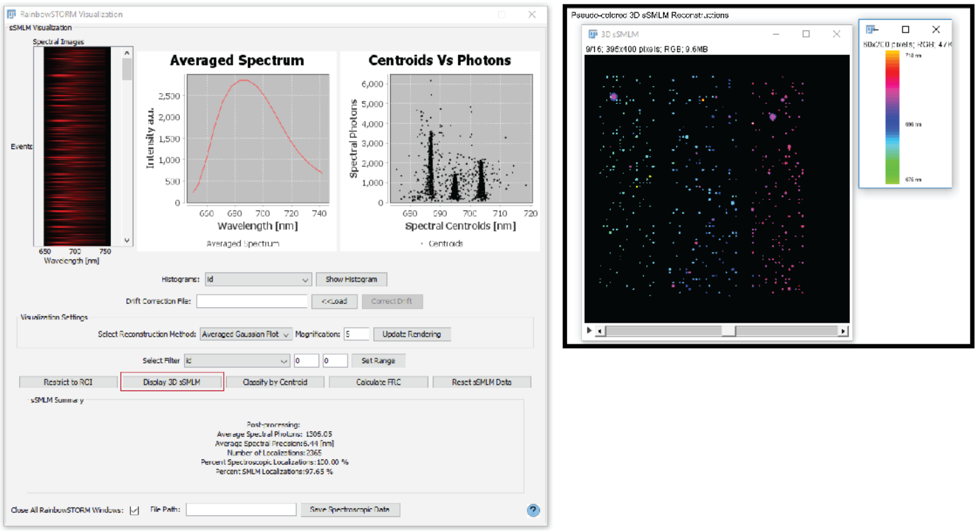

**Figure 30:** Workflow for displaying 3D sSMLM data-acquired recorded using the astigmatism method as a stack of Pseudo-colored super-resolution reconstructions

#### 5.4 Fourier Ring Correlation analysis

Image quality for SMLM image reconstructions can be assessed using the Fourier Ring Correlation (FRC) analysis as described by Nieuwenhuizen (Nieuwenhuizen, et al., 2013), please refer to the referenced publication for full description and optimization methods for FRC analysis. Briefly, FRC analysis takes spatial sampling and localization uncertainty into consideration when determining the resolution of super-resolution reconstructions. To perform this analysis, two images containing localizations from odd and even frames are generated. Next, the Fourier transform for both images are generated. The correlation of the Fourier transforms for both images is calculated over the perimeter of a circle with a given radius in Fourier space and used to generate a plot of the decay of the Fourier correlation with increasing spatial frequency. The Fourier Image REsolution (FIRE) number defined by Nieuwenhuizen et al is the spatial resolution using a FRC threshold of 1/7 and is calculted as the inverse of the spatial frequency at that threshold. The result of the FRC analysis is the FRC plot which also lists the FIRE number (Figure 31).

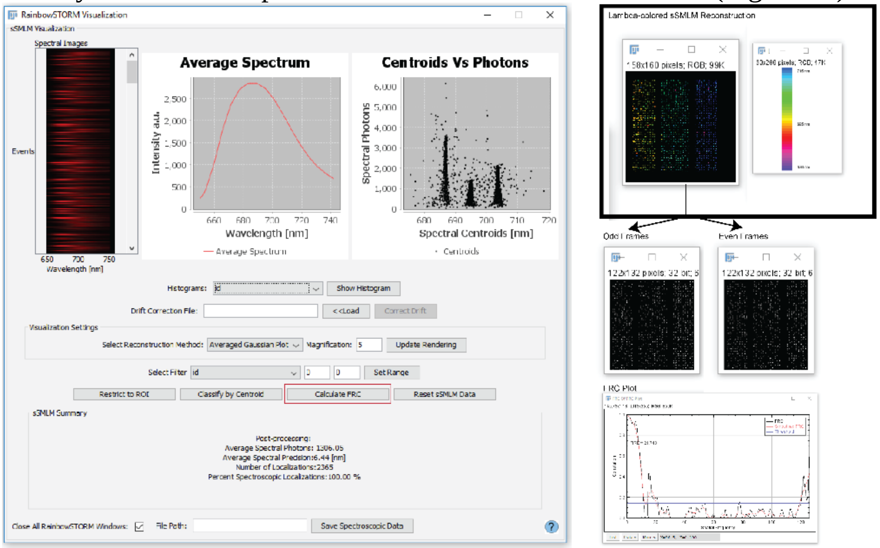

**Figure 31:** Workflow and results for Fourier ring correlation analysis.

### 6. Import sSMLM results

Previously processed sSMLM results can be loaded by selecting the ‘Import sSMLM’ button under the RainbowSTORM menu (Figure 32). Next, the sSMLM results file (ending in ‘_spec.csv’) can be selected and loaded by clicking the ‘Load Results’ button. The camera settings must also be updated to match the settings used to process the original data. The Pseudo-colored images will be reconstructed based on the spatial coordinates of the RainbowSTORM results. Next the ‘Visualize Data’ button can be clicked to launch the visualization screen and Pseudo-colored reconstructions. The results can be updated and saved for later use. The help screen for the import screen can be accessed by pressing the blue question mark button.

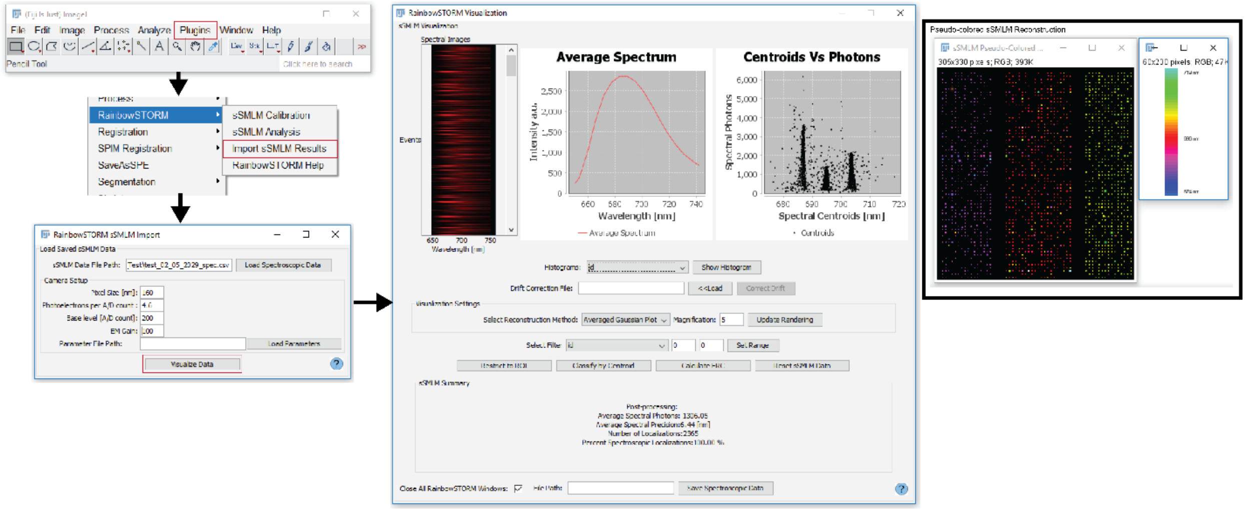

**Figure 32:** Workflow for importing previously saved sSMLM results using the RainbowSTORM import screen

### 7. RainbowSTORM help

The RainbowSTORM help option launches the RainbowSTORM help screen. Figure 33 shows an example of a RainbowSTORM help screen. Additional information, tutorials, the java code as well as this user guide can be found on the RainbowSTORM GitHub page. Please report any issues with the software using the Issues tab on the GitHub page.

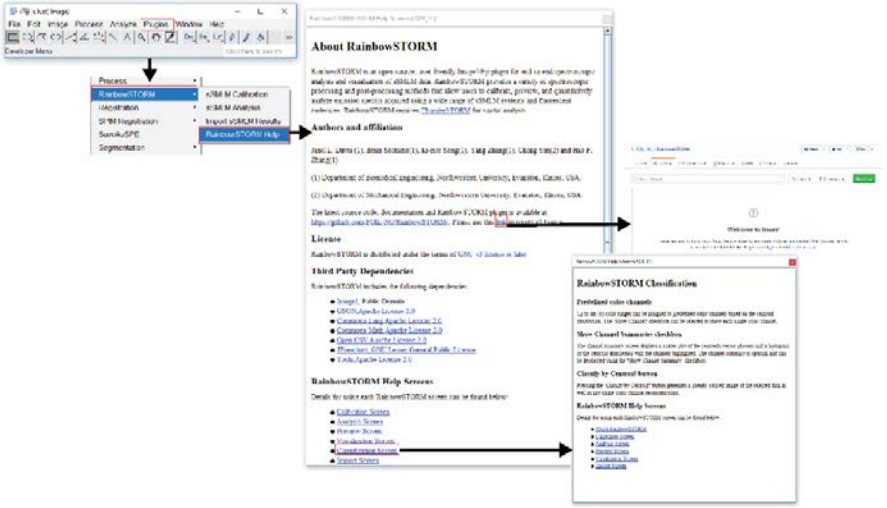

**Figure 33:** Example of using the RainbowSTORM help screen
