## Supplementary Information for "RainbowSTORM: An open-source ImageJ plugin for spectroscopic single-molecule localization microscopy (sSMLM) data analysis and image reconstruction"

**Janel L. Davis[1, †], Brian Soetikno[1, †], Ki-Hee Song[1], Yang Zhang[1], Cheng Sun[2] and Hao F. Zhang[2, *]**

[1]Department of Biomedical Engineering, Northwestern University, Evanston, Illinois, USA.
[2]Department of Mechanical Engineering, Northwestern University, Evanston, Illinois USA

† These authors contributed equally

### RainbowSTORM Algorithms and Equations

The algorithms and equations used for system calibration and spectral analysis of spectroscopic single-molecule localization microscopy (sSMLM) images are detailed in the following sections.

#### System Calibration

The peak pixel positions of known wavelengths from a calibration light source or from multi-color emitter (e.g. Tetraspeck Microsphere, Thermofisher) are selected and fitted using the least-squares method. Grating-based systems are calibrated by fitting the pixel positions to a straight line ($y = a + bx$). Meanwhile, prism-based systems are calibrated by fitting the pixel positions and corresponding wavelength information using either a second-order polynomial ($y = a + bx + cx^2$) or a third-order polynomial ($y = a + bx + cx^2 + dx^3$). The coefficients a, b, c and d represent the offset and the coefficients of the first, second and third orders. The resulting coefficients are used to calibrate the fluorescence emission spectra from single-molecule dyes or fluorescent proteins captured during sSMLM experiments.

#### Global background estimation

An averaging filter is used to estimate the background of the spectral images. Pixel values which are associated with the sample are identified by comparing each pixel value to a threshold. The threshold for this filter can be either user-defined or automatically generated. Figure S1 shows the flowchart of the algorithm used to generate background image (B) from the stack of input images (I). The background image is then subtracted from each frame of the input images to generate a stack of background-subtracted spectral images.

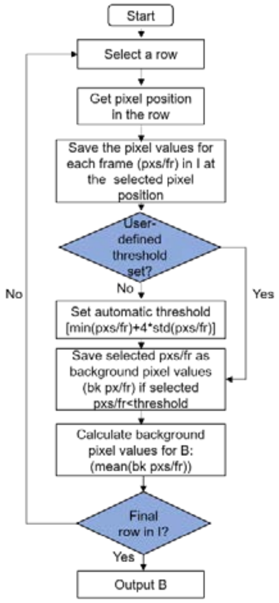

**Figure S1:** Flowchart of the algorithm to generate the background image

#### Overlapping spectra removal

Localizations with spectral images which overlap in space can be excluded from analysis by enabling the 'Remove overlapping spectra' checkbox on the sSMLM analysis module. The localization pixel shift (lps) and the spectral dispersion (sd) are used to remove the overlapping spectra. The lps controls the number of pixels used to capture the point spread function (PSF) of each localization. By default, lps is set to ±1 for a total of three pixels to capture the PSF. The sd captures the number of pixels which cover the spectral image (spectra pixels) and is set based on the sSMLM system. Increasing either the lps or the sd will result in more localizations being removed as overlapping. The thresholds along the x axis is twice the number pixels to capture the spectra and the y axis is twice the number of pixels to capture the PSF. Figure S2 shows the flowchart of the algorithm used to determine whether the spectra are overlapping.

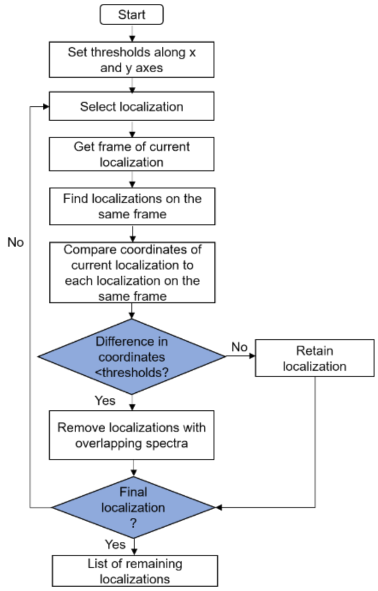

**Figure S2:** Flowchart of the algorithm to remove overlapping spectra

**Spectral centroid calculation**

The intensity-weighted spectral mean or the spectral centroid ($\lambda_{SC}$) of the line spectrum for each localization is calculated by the equation below

$$\lambda_{SC} = \frac{\sum_{\lambda=\lambda_1}^{\lambda=\lambda_n} I(\lambda)*\lambda}{\sum_{\lambda=\lambda_1}^{\lambda=\lambda_n} \lambda} \qquad [1]$$

where $\lambda$ is the wavelength between ($\lambda_1$ and $\lambda_n$) and I is the photon count at that wavelength.

**Spectrum width estimation**

The line spectrum for each blinking event is fit using the least-squares method to a Gaussian function:

$$y = a * e^{-\frac{(x-b)^2}{2c^2}} \qquad [2]$$

The coefficients *a*, *b*, and *c* represent the peak intensity, mean value ($\mu$) and standard deviation ($\sigma$) respectively.

**Spectral precision assessment**

The spectral precision ($\sigma_\lambda$) of each spectrum is estimated as reported by Song et al(Song, et al., 2018):

$$\sigma_\lambda = \sqrt{n_{bg}^2 + n_s^2 + n_{ro}^2 + \sigma_{sse}^2} \qquad [3]$$

where $n_s$, $n_{bg}$, $n_{ro}$ and $\sigma_{sse}$ are the shot noise, background noise, readout noise and uncertainty of the spectral-shift error respectively.

The background noise of the signal is

$$n_{bg} = \sqrt{\frac{2\left(\frac{1024 B_t \eta \Delta\lambda W_p}{64 a \Delta\lambda} * a^3 * \sigma_{PSF}\right)}{3\Delta\lambda W_p (\eta I)^2}} \qquad [4]$$

where $B_t$ is the total background of the spectral signal, $\eta$ is the quantum efficiency , Δλ is the spectral dispersion, $W_p$ is the pixel size, $a = \sqrt{\sigma_{SPE}^2 + \frac{\Delta\lambda^2}{12}}$ where $\sigma_{SPE}$ is the standard deviation of the fitted spectrum, $\sigma_{PSF}$ is the standard deviation of the fitted point spread function and I is the total photon count of the spectrum.

The shot noise of the signal is calculated by

$$n_s = \sqrt{\frac{2\left(\sigma_{SPE}^2 + \frac{\Delta\lambda^2}{12}\right)}{\eta I}} \qquad [5]$$

The readout noise is calculated by

$$n_{ro} = \sqrt{\frac{1024 \sigma_{SPE}^3 \Delta\lambda^2 N_r^2}{3\Delta\lambda W_p (\eta I)^2}} \qquad [6]$$

where $N_r$ is the readout noise per pixel.

The uncertainty of the spectral-shift error limits the spectral precision which can be achieved and can be quantitatively expressed as

$$\sigma_{sse} = \sqrt{\frac{\Delta\lambda^2}{12}} \qquad [7]$$

Figure S3 shows the comparison of the expected spectral precision and the spectral precision estimated with RainbowSTORM using simulated sSMLM images of single-molecule emission events with increasing photon counts. For the estimation, we used Δλ of 5.9 nm/pixel, $\sigma_{SPE}$ of 22 nm, $\sigma_{PSF}$ of 117 nm, $N_r$ of 1 e-, $W_p$ of 16 µm, and $\eta$ of 1.

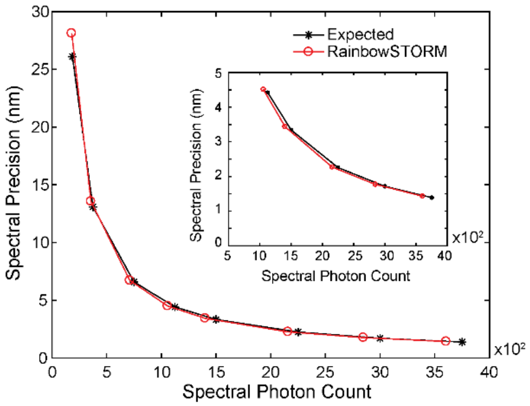

**Figure S3:** Comparison of the expected spectral precision and the average spectral precision estimated by RainbowSTORM at different spectral photon counts [180 to 3800]. Inset shows the zoomed in comparison for the spectral photon count from 1000 to 3800.

### Three-dimensional sSMLM images

Three-dimensional (3D) sSMLM images can be acquired using the astigmatism method (Huang, et al., 2008) by placing a cylindrical lens in the spatial imaging path(Zhang, et al., 2015) as shown in Figure S4.

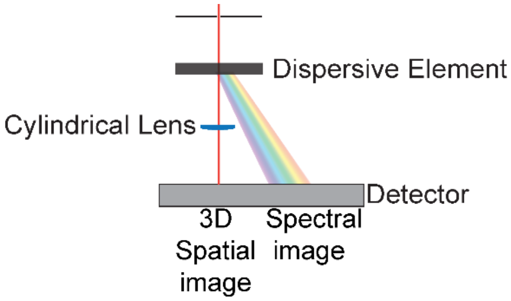

**Figure S4:** General sSMLM system schematic for 3D imaging using the astigmatism method.

### Methods

### sSMLM Optical Setup

The sSMLM system is setup as previously described (Zhang, et al., 2019) and briefly described as follows. A 642-nm CW laser was focused on the back focal plane of a Nikon Ti microscope body and illuminated through a 100× Total Internal Reflection Fluorescence (TIRF) objective lens and a numerical aperture (NA) of 1.49 (CFI Apochromat, Nikon). Fluorescence from the sample is collected by the objective lens and passed through a tube lens then directed to an entrance slit using a mirror. The entrance slit is used to confine the field of view (FOV) of the spatial image to one section of the electron multiply charge coupled device (EMCCD). The photons then pass through a transmission grating (100 grooves/mm, STAR100 Paton Hawksley Education Ltd) which splits the photons into spatial and spectral images with a ratio of ~ 1:3. The remaining photons are then passed through an imaging lens and then focused onto the EMCCD (ProEM, Princeton Instrument) by a second imaging lens.

**Sample Preparation**

COS-7 cells (ATCC) were grown in Dulbecco's Modified Eagle Media (Gibco/Life Technologies) supplemented with 2-mM L-glutamine (Gibco/Life Technologies), 10% fetal bovine serum (Gibco/Life Technologies), and 1% penicillin/streptomycin (10,000 U $mL^{-1}$, Gibco/Life Technologies) at 37°C with 5% $CO_2$. The cells were plated on No. 1 borosilicate bottom eight-well Lab-Tek Chambered coverglass with 30%-50% confluency. The cells were fixed after 48 h in pre-warmed 3% paraformaldehyde and 0.1% glutaraldehyde in phosphate-buffered saline (PBS) for 10 min, washed with PBS twice and quenched with freshly prepared 0.1% sodium borohydride in PBS for 7 mins. The cells were then rinsed three times in PBS at 25°C. The fixed cells were then permeabilized with blocking buffer (3% bovine serum albumin (BSA), 0.5% Triton X-100 in PBS) for 20 min and then incubated with primary antibodies (sheep anti-tubulin, 10 ug $mL^{-1}$, rabbit anti-PMP70 and mouse anti-TOM-20, 5 ug $mL^{-1}$) in blocking buffer for 1 hr at room temperature and rinsed with washing buffer (0.2% BSA, 0.1% Triton X-100 in PBS) three times. The cells were then incubated in donkey secondary antibody conjugates (anti-sheep Alexa Fluor 647, anti-mouse CF660C and anti-rabbit CF680) in blocking buffer for 40 min and then washed three times with PBS and stored at 4°C. The secondary antibodies were prepared as previously described and briefly described here. 0.5 uL of 5mM dye (NHS-ester functionalized Alexa Fluor 647 (AF647, Thermofisher), CF660C and CF680 (Biotinum)) were combined at 25°C with 100 uL of 1 mg $mL^{-1}$ IgG/IgY antibodies in PBS and sodium bicarbonate (10 uL of 1 M). The mixture was incubated overnight then purified by a Nap-5 size exclusion column. The concentrated sample was then extracted using an Amicon Ultra-0.5 Centrifugal Filter unit to give 1-2 dyes per antibody. Finally, the absorption and emission spectra of the dyes were tested using a NanoDrop Spectrophotometer and then stored at 4°C.

**sSMLM Image Acquisition**

**System Calibration**

Using a calibration light source (Neon lamp, 6032 Newport) reference images were captured by using a narrow slit. The images in the spatial domain was a narrow straight line while the spectral images showed multiple spectral lines representing known emission maxima related (640.23 nm,703.24 nm,724.52 nm, and 743.89 nm) to the calibration light source.

**Experimental sSMLM Data**

Imaging buffer (50 mM Tris (pH=8.0), 10 mM NaCl, 0.5 mg $mL^{-1}$ glucose oxidase (Sigma, G2133), 2000 Uq/mL catalase (Sigma, C30), 10% (w/v) D-glucose, and 100 mM cysteamine) was added to the COS-7 cells and the cells were imaged using the sSMLM system described above. The exposure time was set to 10 ms, pixel size of the system was 160 nm, the analog to digital unit (ADU) was 4.6 $e^-$/analog to digital count, and the electron multiplying gain (EM Gain) was 100. For this experiment 30000 frames were recorded.

**Simulated sSMLM Data**

We simulated 500 frames of sSMLM images with a back-projected pixel size of 160 nm, camera base level of 200 digital counts, EM Gain of 100, and ADU of 4.6 $e^-$/analog to digital count. For spectral precision testing, the simulated emitter was AF647 with total photon counts of

250,500,1000,1500, 2000, 3000, 4000 and 5000. The photons from the simulated emitters were split into spatial and spectral images using a 1:3 ratio. The spatial image of each single-molecule event was modeled as a 2D gaussian with a mean of 0 and std of 0.73 pixels. A reference line spectrum for AF647 was convolved with the point spread function image to generate the spectral image. The shifted spectral image was set based on a calibration lamp with known emission maxima (485.5 nm, 546.5 nm, 611.6 nm, and 707 nm). The locations of each localization event and its associated spectrum were randomly assigned. For multicolor simulated data reference spectra of CF660 and CF680 were used to generate spectral images.
